# Global elongation and high shape flexibility as an evolutionary hypothesis of accommodating mammalian brains into skulls

**DOI:** 10.1101/2020.12.06.410928

**Authors:** Vera Weisbecker, Timothy Rowe, Stephen Wroe, Thomas E. Macrini, Kathleen L. S. Garland, Kenny J. Travouillon, Karen Black, Michael Archer, Suzanne J. Hand, Jeri Berlin, Robin M.D. Beck, Sandrine Ladevèze, Alana C. Sharp, Karine Mardon, Emma Sherratt

**Author notes:** Corresponding author e-mail addresses.

## Abstract

Little is known about how the large brains of mammals are accommodated into the dazzling diversity of their skulls. It has been suggested that brain shape is influenced by relative brain size, that it evolves or develops according to extrinsic or intrinsic mechanical constraints, and that its shape can provide insights into its proportions and function. Here, we characterise the shape variation among 84 marsupial cranial endocasts of 57 species including fossils, using 3D geometric morphometrics and virtual dissections. Statistical shape analysis revealed four main patterns: over half of endocast shape variation ranges between elongate and straight to globular and inclined; little allometric variation with respect to centroid size, and none for relative volume; no association between locomotion and endocast shape; limited association between endocast shape and previously published histological cortex volumes. Fossil species tend to have smaller cerebral hemispheres. We find divergent endocast shapes in closely related species and within species, and diverse morphologies superimposed over the main variation. An evolutionarily and individually malleable brain with a fundamental tendency to arrange into a spectrum of elongate-to-globular shapes – possibly mostly independent of brain function - may explain the accommodation of brains within the enormous diversity of mammalian skull form.

## Introduction

The origin of mammals is demarcated by two key innovations: the evolution of a large brain (Rowe et al. 2011), and a fundamental reorganisation of the skull and particularly the endocranial cavity housing the brain (Maier 1993; Koyabu et al. 2014). Fitting the brain into the evolving skull with its diverse functions requires tight evolutionary and developmental integration between the two (Hanken and Thorogood 1993; Richtsmeier and Flaherty 2013). However, the evolution of this relationship – which can be characterised through studies of the endocranial cavity, or “endocast” (Jerison 1973; Macrini et al. 2007a) - is little understood on larger evolutionary scales, with most research focused on the large-brained primates (Bienvenu et al. 2011; Aristide et al. 2016; Zollikofer et al. 2017) or experimental rodent models (Lieberman et al. 2008; Nieman et al. 2012), and focused on the understanding of the exceedingly large human brain.

The most widely discussed hypotheses of primate brain shape evolution – mostly done on cranial endocasts representing the bony braincase cavity mostly occupied by the brain - focus on allometry (change with size) either related to absolute or relative brain sizes. For example, the influential “spatial packing” hypothesis posits that relatively larger brains can evolve to be flexed at the base and rounded, because this packs the brain most efficiently into a limited space (Biegert 1957; Lieberman et al. 2008; Bastir et al. 2011; Zollikofer et al. 2017). “Spatial packing” is thought to explain the coevolution of mammalian brain and cranial base shape in primates (Gould 1977; Ross and Ravosa 1993; Ross and Henneberg 1995; Bastir et al. 2011) and mouse strains (Lieberman et al. 2008). There is also general support that absolute brain size (rather than relative brain size) correlates with primate endocast shape, probably differentially in different clades (Aristide et al. 2016; Sansalone et al. 2020). However, beyond primates, there are multiple hypotheses of how the mammalian brain is shaped, involving different levels of organisation. For example, the growing mammalian brain appears to adapt to endocranial shape changes (Jeffery and Spoor 2006; Macrini et al. 2007c; Budday et al. 2015); physical internal constraints of neuronal connectivity determine the shape at least of the neocortex (Atkinson et al. 2015; Mota and Herculano-Houzel 2015); and pleiotropic genetic effects appear to shape both cranial vault and brain and may limit the availability of endocast shapes to selection (Hanken and Thorogood 1993; Koyabu et al. 2014). Lastly, individual brains can grow and shrink substantially in the lifetime of an individual, expanding the osseous braincase (Dechmann et al. 2017), but there is also evolutionary (Koyabu et al. 2014) and developmental (Nieman et al. 2012; Richtsmeier and Flaherty 2013) evidence that the dorsal cranial vault adapts to increases in brain size, not vice versa.

Characterizing the patterns of mammalian brain shape evolution represents a necessary first step in unravelling the complex intrinsic and extrinsic determinants of mammalian brain shape and development. Metatherian mammals (marsupials and their closest extinct relatives) are an ideal study group for this: they are an old (> 110 million years; Eldridge et al. 2019), monophyletic, phylogenetically well-understood radiation of mammals and occupy all terrestrial ecological niches except for active flight. Body masses of living marsupials span three orders of magnitude (2.6 g up to 85 kg; Weisbecker et al. 2019), with extinct representatives weighing up to 2.7 tonnes (Wroe et al. 2004), making them particularly well-suited to resolve current debates on whether brain size and brain shape are correlated. In addition, the development of living marsupials makes them a more suitable radiation for studies of brain morphology: marsupials are born at highly altricial stages compared to placentals, after maximally 30 days of pregnancy, and grow during uniform three-phase lactation period (Hinds 1988), in which the milk composition is tailored to the requirements of the pouch young. Marsupial brain development occurs nearly entirely *ex utero* during lactation (Smith 1997). Reproductive traits relating to maternal investment are an important influence on the variability of brain growth (e.g. Bennett and Harvey 1985; Barton and Capellini 2011). The relative uniformity of marsupial reproduction provides an additional argument for this clade to be a test case for mammalian brain evolution (Pirlot 1981; Weisbecker and Goswami 2010, 2011b; Weisbecker et al. 2015), particularly because marsupial and placental brains are fundamentally similar in structure (Ashwell 2010), relative brain size (Weisbecker and Goswami 2010), and neuronal scaling (Dos Santos et al. 2017).

Here we use geometric morphometrics to analyse three-dimensional endocast shape in a sample of 57 marsupial species across all major clades, including 12 fossil species. We aim to determine if a common pattern of shape variation emerges in marsupials, and whether suggestions for allometric patterning of endocast shape, posited for primates, can be generalised to marsupials and could generally explain mammalian brain shape evolution. We also assess suggestions that endocast shape corresponds with locomotor function (Ahrens 2014; Bertrand et al. 2019b) and phylogeny (e.g. Silcox et al. 2009; Thiery and Ducrocq 2015; Bertrand et al. 2016; Bertrand et al. 2019b). To further understand if brain shape is indicative of internal brain structure, we use volume dissections of our endocasts and a limited dataset of neocortical and isocortical grey matter volumes (noting that iso- and neocortex largely refer to the same structure; Palomero-Gallagher and Zilles 2015) to assess the degree to which brain partition volumes and brain shape correspond in a large, phylogenetically diverse mammalian radiation.

## Methods

### Specimens

This study is based on cranial endocasts, which have been shown to be a good approximation of brain shape for mammals and marsupials in particular (Jerison 1973; Macrini et al. 2007a). Virtual reconstructions of endocasts from conventional and micro-Computed Tomography (μCT) scans from adult skulls of 57 marsupial species were prepared in Materialise Mimics (v.17-20) through virtually “flood-filling” the endocranial cavity of specimens, mostly by two operators, with some help from technical staff. The list of specimens with their museum numbers is available in the main data file used in this analysis (Supplementary Table 1). The 3D endocasts files that were landmarked are available on Figshare (10.6084/m9.figshare.12284456; a selection of endocasts is illustrated in Fig. 1)

**Figure 1:**
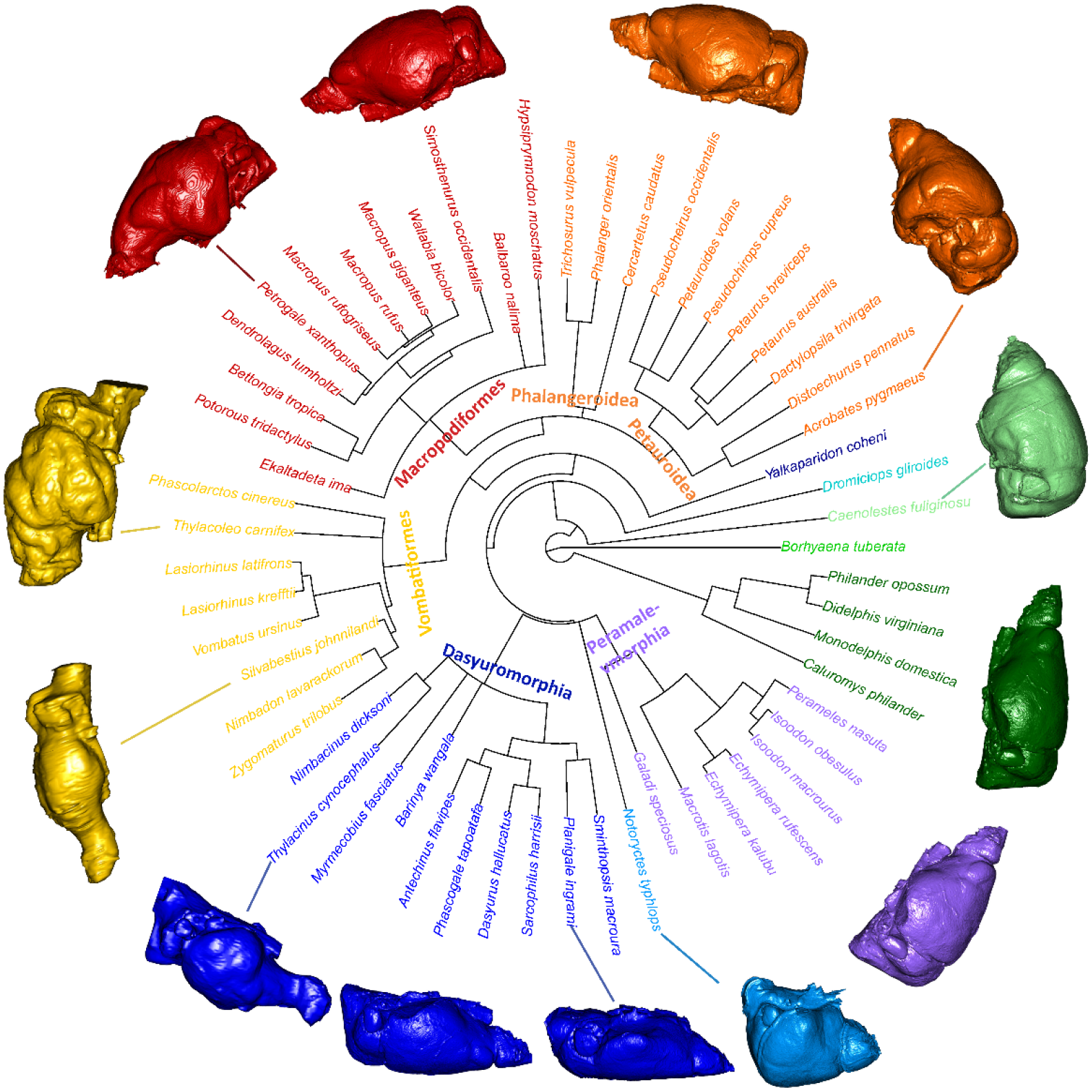
A summary of the species and one of the three phylogenetic topologies used, with the “main clade” designations and their colour scheme used throughout the figures of this study. Crosses next to species names denote fossil species. Endocasts arranged radially reflect the mean shapes for each clade plus some notable endocasts of individual species (connected to their species name by lines) to highlight the shape diversity.

One specimen per species was sourced for 38 species, and multiple specimens, where possible including males and females, were sourced for 19 species to assess how representative a single endocast is of a species. Scans of extant species were conducted at either the University of Texas High-Resolution X-ray Computed Tomography Facility (Austin, Texas, USA), a SkyScan 1074 at the University of Queensland (School of Biological Sciences), or a Siemens Inveon Pet-CT scanner at the UQ Centre of Advanced Imaging at the University of Queensland (Australia). To allow easy manipulation of large scans, these were cropped to just the brain case and/or subsampled by either using every second slice only and/or reducing the resolution of each slice in ImageJ (Schneider et al. 2012). All but three endocasts were derived from museum specimens (for accession numbers, see Supplementary Table 1). The study also used post-mortem CT scans of one captive dunnart (*Sminthopsis macroura*), and two captive koalas (*Phascolarctos cinereus*), whose deaths were unrelated to this study. The *S. macroura* specimen was obtained in 2008 from Melbourne University under Animal Ethics MAEC (Vic) License No. 06118. The koalas were scanned under University of Queensland permit SVS/405/12. CT scans of twelve fossils were also included. Note that the Tasmanian tiger (*Thylacinus cynocephalus*), which went extinct in 1936, is here assigned “extant” status because of its very recent extinction. Small areas of incomplete surfaces at the rear of the cerebellum were virtually repaired in *Dromiciops gliroides* and *Petaurus australis*. This involved the scaling and manual fitting of similarly curved cerebral surfaces to complete the missing surface. *Caenolestes fuliginosus* was scanned in two sessions so that two scans were reconstructed and the resulting two endocasts were retrospectively fit together, using the global registration tool in Mimics.

### Endocast landmarking and protocol comparisons

Endocast shape was characterised in three dimensions with 29 fixed landmarks, ten curve semilandmarks, and 32 surface sliding semilandmarks (Figure 2; for anatomical description and numbering of the landmarking protocols, see Supplementary Information 1). The landmarking of endocasts represents a challenge because the cerebral hemispheres have large areas with few landmarkable homologous features. This was exacerbated due to varying scan resolutions, and the high diversity of brain shapes within our sample, so that landmark repeatability was expected to be an issue (Pereira-Pedro and Bruner 2018). To capture endocast shape effectively, and test the repeatability of our resulting landmark dataset, one of us (VW) landmarked each endocast twice and trialled two different protocols for surface landmark placement. This included a fully manual protocol using only the Checkpoint programme (Wiley 2015). The other protocol used the Checkpoint software (Wiley 2015) for fixed and curve semilandmarks, and automatic placement of surface semilandmarks using the *Morpho* R package (v. 2.8).

**Figure 2:**
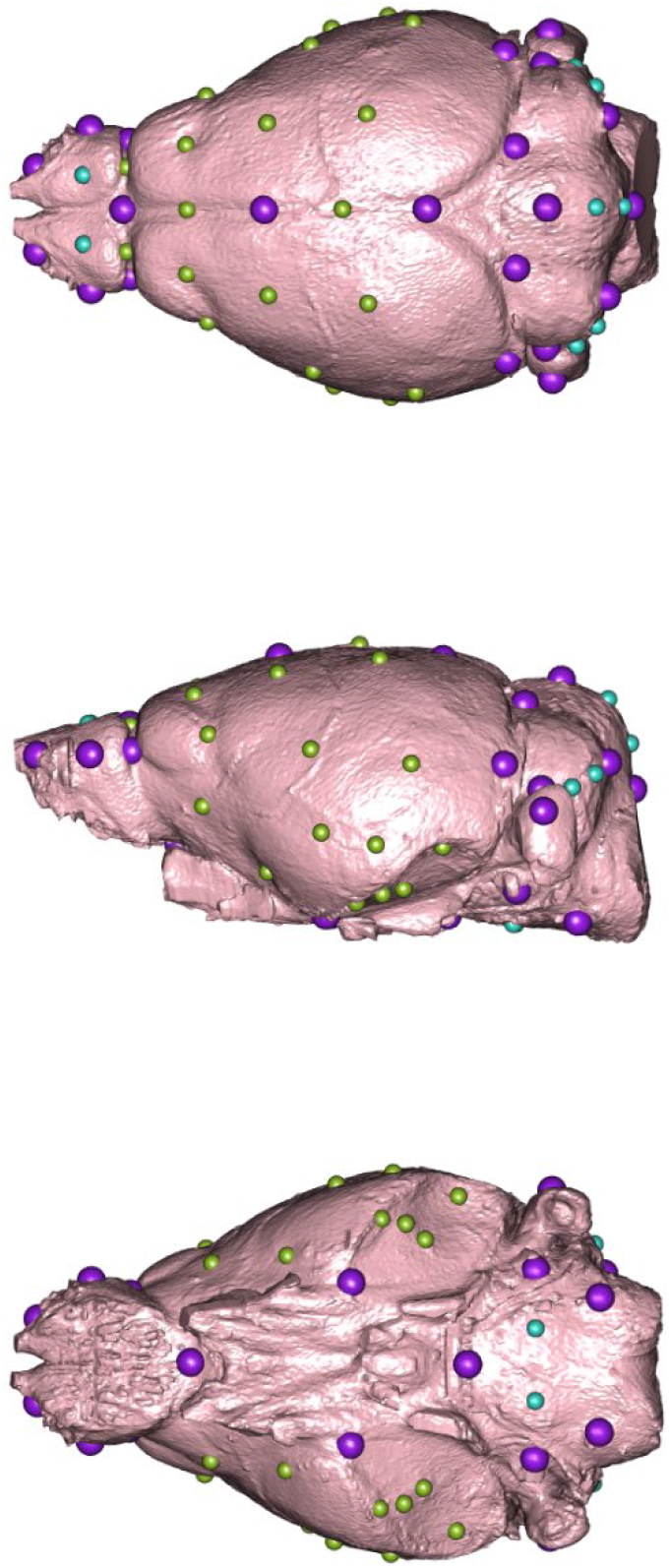
Landmark placement on the brain of *Phascogale tapoatafa* JM12395 (the specimen used to produce the atlas for automatic landmark placements), warped to the mean shape in geomorph. Large, purple spheres are fixed landmarks; turquoise spheres are curve semilandmarks; and green spheres are patch semilandmarks.

The performance and similarity of the two surface landmarking protocols was tested in several ways. To assess repeatability of landmarking, the shape variance from each replicate and each protocol was evaluated against among-species variance using a Procrustes analysis of variance (ANOVA) (Zelditch et al. 2012). To test the correspondence between the two protocols, principal components analyses (PCA) was used to construct two multi-dimensional spaces, and the distribution of species in each space was compared using a Mantel test in the *Vegan* R package v. 2.5-6 (Oksanen et al. 2016). In addition, scatter plots of the first two principal components for each protocol, as well as graphs of the shape changes along the axes, were produced for visual comparison.

All automated landmarking procedures, analyses and results were conducted in the R statistical environment (R core team 2018) and can be replicated with our code on github; the ply files that can be used for automatic landmarking are in 10.6084/m9.figshare.12253409.

### Endocast partitions and additional neo/isocortex data

The endocasts were virtually “dissected” into four partitions representing the cerebrum, olfactory bulbs, cerebellum, and brain base (the protocol for how the boundaries were defined is in Supplementary information 1). Volumes of the four dissected partitions were measured using Mimics (versions 17-20) and averaged per species if multiple specimens per species were available. Such partitions are only approximate representations of brain regions because they do not allow for internal brain boundaries. However, this method has been successfully used as a general indicator of overall brain proportions (Sakai et al. 2016; Sakai et al. 2018; Bertrand et al. 2019a) and is used with this caveat here. A log-shape ratio approach was used to standardise the partition volumes with respect to scale (Mosimann 1970). This was done by deriving the logarithm of the value obtained by dividing each partition volume by the geometric mean of all volumes for a specimen, which removes the isometric component of size and is thus comparable to the scaling of landmark configurations during generalised Procrustes analysis.

Because the neocortex comprises a large part of overall volume of the marsupial brain (Pirlot 1981) and mammalian cerebral hemispheres (Jerison 2012), we also assessed if any of the endocast shape or cerebral hemisphere volume variation in our sample was suggestive of neocorticalisation – where some marsupials, particularly diprotodontians (Pirlot 1981), or extinct mammals in general (Jerison 2012) have relatively larger or smaller neocortices, respectively. While a confident dissection of the neocortical portion of the cerebral hemispheres was not possible, we matched data from two publications with a sample that was overlapping ours. We computed the log-shape ratios of neocortex volumes from Pirlot (1981) for 17 species of marsupials, including members of Didelphimorphia, Dasyuromorphia, Peramelemorphia and Diprotodontia; and isocortex grey matter volumes for 19 Diprotodontian species from Jyothilakshmi et al. (2020). These log-shape ratios were based on the larger datasets of histology-based brain region volumes of which the neo/isocortical volume data were part (for computation of these from the full datasets, refer to the relevant file on github). In one case for Pirlot (1981) and five cases for Jyothilakshmi et al. (2020), a species in our dataset was matched with a value from a congeneric species of a similar body weight (noted in Supplementary Table 1).

### Phylogeny, body mass, and locomotor mode

The details on tree estimation are in Supplementary information 1; the tree files are part of the raw data folder on this study’s github repository. For all phylogenetic analyses, we used three topologies differing in their placement for the fossil *Yalkaparidon*, whose phylogenetic affinities are uncertain (Beck et al. 2014). Figure 1 displays one of these phylogenetic topologies. All polytomies were considered as soft polytomies, and randomly resolved to zero branch lengths using the multi2di function from the *ape* package v.5..4-1 (Paradis et al. 2004) in R. In all phylogenetically-informed analyses in this paper, the use of different topologies had minimal impact on test statistics and no impact on significance levels, and thus an average of the test statistics from the three trees is presented here.

Body mass estimates for extant species were taken from Weisbecker et al. (2013), and for fossil species were derived from multiple publications as outlined in Supplementary Table 1 (Wroe et al. 1999; Argot 2003; Turney et al. 2008; Travouillon et al. 2009; Black et al. 2012; Sharp 2016).

All species were allocated broad locomotor categories: arboreal, terrestrial, gliding, scansorial, hopping and fossorial (Supplementary Table 1). Locomotor data for Australian species were taken from Weisbecker et al. (2020), and for American species was taken from Nowak (2018). Note that the locomotion of the tree kangaroo *Dendrolagus lumholtzi* was scored as “hopping”, because this is the main locomotor mode of the species despite the fact that it lives in trees. Note also that only two dasyurids (the yellow-footed antechinus and the northern quoll) in the dataset were classified as “scansorial”; due to this small sample size, these two were classified as “arboreal” because most other dasyurids are terrestrial, so that a capacity to climb as part of the scansorial locomotor repertoire is likely to have arisen from terrestrial ancestry. The only fossorial species sampled, *Notoryctes typhlops*, was excluded from the locomotor analyses.

### Shape analyses

Landmark coordinates were subjected to a generalized Procrustes superimposition and projection into tangent space (Rohlf and Slice 1990) using the R package *geomorph* v.3.3.1 (Adams et al. 2020). The semilandmarks identified as curves were permitted to slide along their tangent directions in order to minimize Procrustes distance between specimens (Gunz et al. 2005); semilandmarks placed as patches were allowed to slide in any direction on two planes. The resulting Procrustes shape coordinates were averaged over the two replicates per specimen. We then performed a second Procrustes fit on the averaged shape coordinates to account for object symmetry (Klingenberg et al. 2002). Only the symmetric component of shape was used as shape variables in the subsequent analyses.

We performed a PCA of species-averaged shape variables to investigate whether variation was restricted to one main axis of brain shape variation, or whether variation was diffuse across several PC axes. For this we examined and plotted all PCs explaining more than 10% of variation. We estimated how much influence allometry has on the primary axis (PC1) by using phylogenetic generalized least squares (PGLS) analysis and evaluating a model of PC1 scores against log centroid size. Note that we used the multivariate generalisation of PGLS (Adams 2014b) implemented in procD.pgls function of *geomorph* for bivariate and multivariate analyses. To visualise the shape changes associated with each PC axis, we used a surface warping approach (Sherratt et al. 2014) whereby an average-shaped triangular mesh is warped to the shape at the minima and maxima of each PC axis using thin-plate spline. The average-shaped mesh was created by taking a species’ endocast the shape of which was most similar to the hypothetical mean shape of all species (*Phascogale tapoatafa* JM12395) and warping the mesh to this hypothetical mean shape using the plotRefToTarget function in geomorph. To aid interpretation of the surface warps, we also produced landmark displacement graphs depicting the direction and magnitude of shape change vectors of each landmark from the mean shape to the PC minima and maxima. To visually depict the diversity of endocast shapes for each major clade as colour-coded in Figure 1, we also warped the mean shape mesh to the mean shape configurations for each of seven clades that had multiple species. We also display some of the more unusual endocasts discussed in this manuscript (Figure 1).

### Phylogenetic signal, allometry, and locomotor mode

To assess the amount of endocast shape variation that can be explained by phylogenetic relatedness, we computed K_mult_, an adaptation of Blomberg’s K that identifies phylogenetic signal in a multivariate context (Adams 2014a). K_mult_ was computed for endocast shape, centroid size, and brain region volume (details below) to test the degree to which variation in brain shape, size and volumetric proportions correspond to Brownian-motion evolutionary processes.

To assess the evolutionary allometry in endocast shape, body mass and centroid size were considered. Centroid sizes were calculated from the endocast landmarks, and we also assessed to see how well it correlated with endocast volume. We performed a PGLS analysis of the shape variables against log-transformed endocast size, and statistical assessment was done by permutation, using 1000 iterations for each model. We also calculated a measure of relative brain size, using the residuals of a linear regression of log-centroid size against log-body mass, which in our sample represents a good approximation of brain/body mass scaling in marsupials (Weisbecker and Goswami 2011a). To visualise how endocast shape is predicted by size, we calculated the regression score (Drake and Klingenberg 2008), which is a univariate summary of the direction of the multivariate regression vector (calculated during the PGLS), and produced landmark displacement graphs of endocast shape for small and large brain sizes.

To assess if endocast shape was predicted by locomotor mode, we used PGLS to evaluate a model of shape variables against log-centroid size plus a factor of locomotor categories (details above). We included analyses to discount the possibility that this relationship might be influenced by an interaction or difference in intercept between size and locomotor mode.

### Intraspecific variation of endocast shape

To assess whether within-species diversity of endocast shapes might have an important impact on the results of our previous between-species analyses, we compared among-species brain shape variation to variation within species. To do this, we conducted a PCA of shape variables of all specimens, calculated the Procrustes distances between all pairs of specimens, and plotted frequency distributions of the pairwise distances separated into within-species and among species groups.

### Comparison of endocast shape with partition volume evolution, and relationship with histological neo/isocortical volumes

For an estimate of how well endocast partition volumes correspond with their shapes, the distribution of species in multi-dimensional PC space between the shape *vs*. volume PCA was computed using a Mantel test in the *Vegan* package (Oksanen et al. 2016) in R. We also computed phylogenetic signal (K_mult_) and evolutionary allometry of brain partition volumes for comparison with K_mult_ of shape variation. Lastly, we assessed if there was any association between shape, cerebral volume, and the literature-derived neocortical/isocortical data (explained above), which might be the case if changes in the relative size of the neocortex are reflected in changes in the relative size of cerebral hemisphere volume. Such an association would suggest that brain shape or volume can be used as an indicator of neocorticalisation, where the neocortical part of the cerebrum expands disproportionately (Jerison 2012; Bertrand et al. 2019a). These analyses were done by evaluating PGLS models of the association between cerebral log-shape ratio and PC2 and neocortex/isocortex grey matter volumes.

## Results

### Endocast landmarking protocol comparisons

The manual and automated placement protocols for surface semilandmarks performed nearly equally well on several analyses: in the Procrustes ANOVA comparing replicates, the automated protocol had slightly higher repeatability (0.91 with manual *vs*. 0.92 with automated placement). The Mantel test of PC score matrices revealed a very similar arrangement of species in PC space (matrix correlation = 0.97), and shape variation along PC1 was near-identical as well (Supplementary Fig. 1). Due to the slightly higher repeatability and better replicability of the automated placement, we present the results from this protocol; however, all analyses can be re-run using the manual protocol in our github code, with near-identical results.

### Endocast shape

The main variation in endocast shape is heavily concentrated on PC1 (Fig. 3), accounting for more than half (56%) of overall brain shape, with PC2 accounting for an additional 12.5%; the subsequent PCs explain 7% or less of the overall variation and are not considered here.

**Figure 3:**
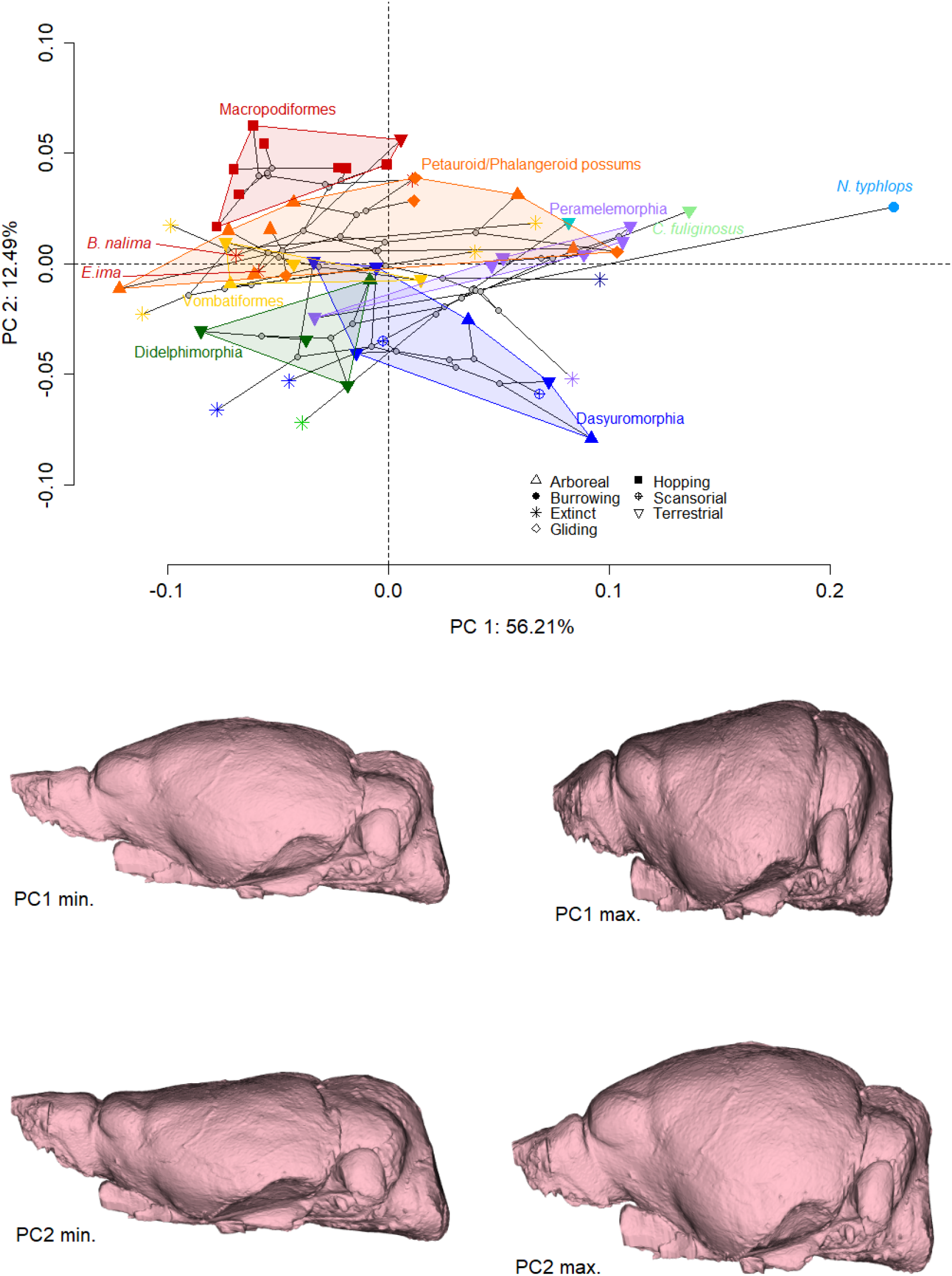
Phylomorphospace (with one of the phylogenies superimposed over the scatter plot) of principal component 1 vs PC2 for 57 species of marsupials according to their brain shape (top) and corresponding shape variation illustrated as warped endocasts representing the highest and lowest PC1 and 2 scores (bottom). Polygons are drawn around living members of all major clades. Stars represent fossil species.

The shape variation along PC1 is almost entirely related to anteroposterior stretching/compression that results in cylindrical, slender shapes at low scores, and shorter, wider, and deeper (and hence more compact and rounded) shapes at high scores (Fig. 3; landmark displacement graphs in Supplementary Fig. 2). Thus, more “stretched” brain shapes at low PC1 scores have longer, narrower olfactory bulbs, cerebra, cerebella, and brain bases. High-PC1 brains also tend to have steeper inclines of the ventral cerebral hemispheres relative to the brain base (Fig. 3, see also supplementary movie 1 (10.6084/m9.figshare.13331273) and landmark displacement graphs in Supplementary Fig. 2).

We also ascertained that the shape variation along PC1 is not driven by two very round-brained outliers, namely the notoryctimorphian *Notoryctes typhlops* (the highly derived marsupial mole) and the paucituberculatan *Caenolestes fuliginosus* (a so-called “shrew opossum”; Fig. 1); removal of these species reduces the magnitude, but not the direction, of shape variation along PC1 and the arrangement of species relative to each other in PC1/2 morphospace remains near-identical (Supplementary Fig. 3).

At first glance, PC2 (accounting for 12% of shape variation) appears to separate diprotodontians from other marsupials. Vombatiforms (wombats, koalas, and fossil relatives) and “possums” (Phalangeroidea and Petauroidea) have overall lower PC2 scores compared to the macropodiform kangaroos (Fig. 3). However, extending each clade’s morphospace to also include fossil species (Supplementary Fig. 4) reveals extensive overlap of macropodiforms, vombatiforms, and petauroid/phalangeroid possums. High scores of PC2 capture a mostly dorsal and slightly posterior expansion of the cerebral hemispheres relative to the olfactory bulbs and the cerebrum, such that the cerebrum appears enlarged, as well as a slight forward shift of the brain base (Fig. 2; Supplementary Fig. 2 and warp movie Supplementary movie 1). Variation in cerebral hemisphere volume is the visually most conspicuous feature of PC2. Notably, four distantly related fossil species are among the eight lowest-scoring species on PC2: *Borhyaena tuberata* (an early Miocene member of Sparassodonta, which is outside the marsupial crown-clade), *Galadi speciosus* (an early Miocene stem-peramelemorphian), *Barinya wangala* (an early-to-middle Miocene dasyuromorphian), and *Nimbacinus dicksoni* (a middle Miocene thylacinid).

### Phylogenetic signal, allometry, and locomotor mode

Overall phylogenetic signal (the degree to which Brownian motion evolution along the phylogeny explains the distribution of values), as measured by K_mult_, was moderate (0.48) for shape but much higher for size (1.02). As also visible in PC2 of the PC space, each clade tended to have its own brain shape, but these shapes are broadly similar compared to the diversity of shapes within each clade (see Fig 1, Supplementary Fig. 5)

Size has a significant but minor association with endocast shape, explaining around 7% of shape variation regardless of whether centroid size (highly-correlated with, and thus equivalent to, brain volume; Table 1) or body mass is used as a size proxy (Table 1), Relative brain size (residuals of the regression of log brain volumes against log body mass) was not significantly associated with brain shape, although this relationship was only just beyond our significance threshold of 0.05 (Table 1). Plotting the regression score against log-centroid size (Fig. 4) revealed that vombatiforms (wombats, koalas, and fossil relatives) and peramelemorphians (bandicoots and bilbies) have brain shapes that do not follow any allometric pattern; removing these two groups from the regression analysis resulted in a higher contribution of evolutionary allometry to brain shape in the remaining marsupials, explaining up to 17% of shape variation (Table 1). A PGLS of PC1 and log centroid size showed that only 12% of PC1 is explained by centroid size (Table 1). However, the variation explained by allometry appears to be a subset of the much greater variation along PC1: the landmark displacement graphs describing the shape change from the predicted values for a small compared to a large endocast (Fig. 4) are very similar to the shape changes associated with minimum and maximum PC1 scores.

**Figure 4:**
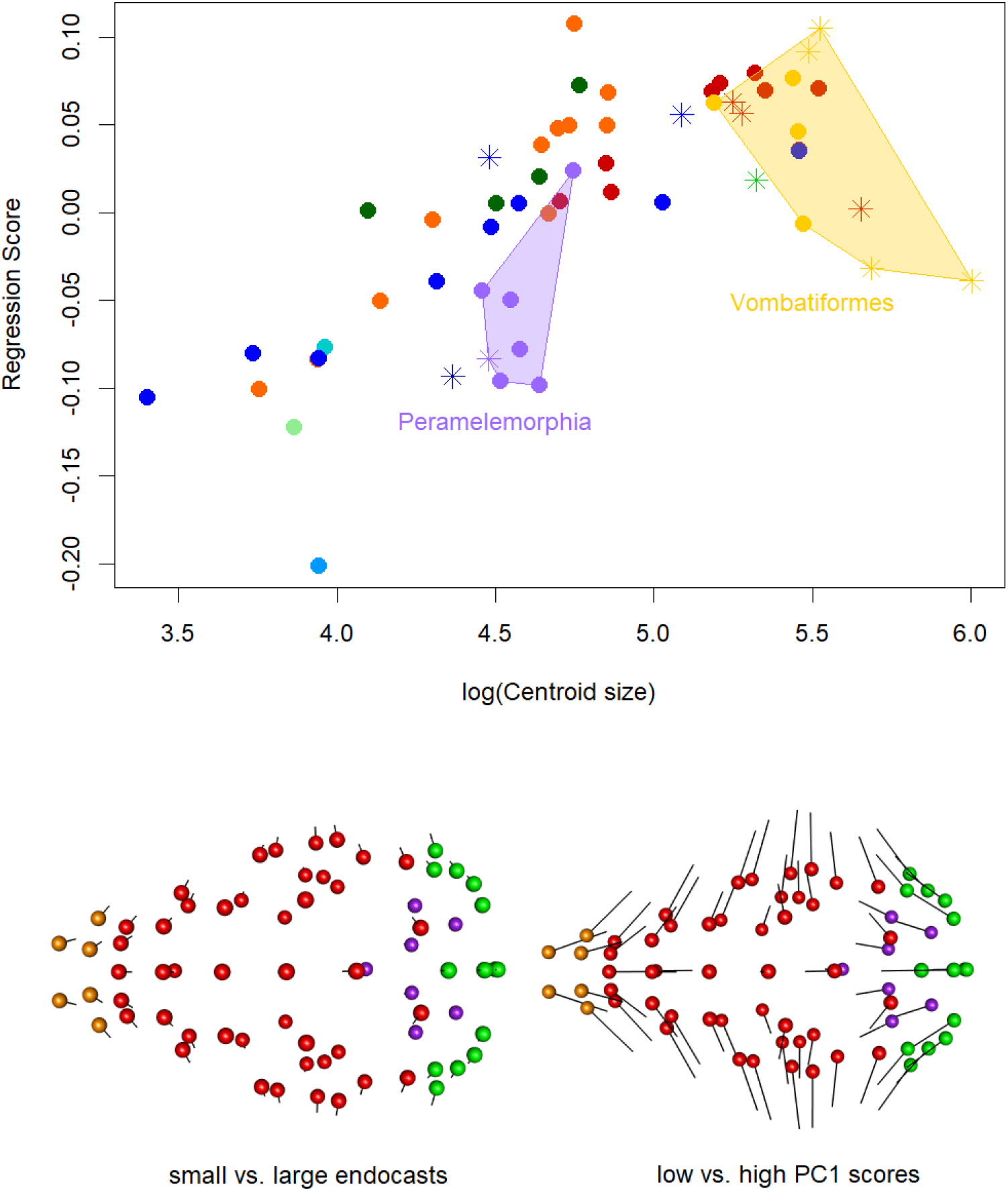
Top: Plot of regression score against log-centroid size showing a small signal of evolutionary allometry in endocast shape (R^2^=0.072, *p*=0.001). Colouration as in Fig. 1, stars are fossil species. Bottom left: Shape changes associated with small to large centroid sizes, shown as landmark displacement graph where ball represents small endocasts and lines point towards large configuration. Bottom right: Shape changes associated with PC1 for comparison with the allometric pattern, where balls indicate the landmark configuration of low PC scores and lines point towards the configuration of shapes with high PC1 scores. Ball colours differentiate landmarks over the surface of four brain regions: olfactory bulbs (orange), cerebrum (red), cerebellum (green), and brain base (purple)

**Table 1:**
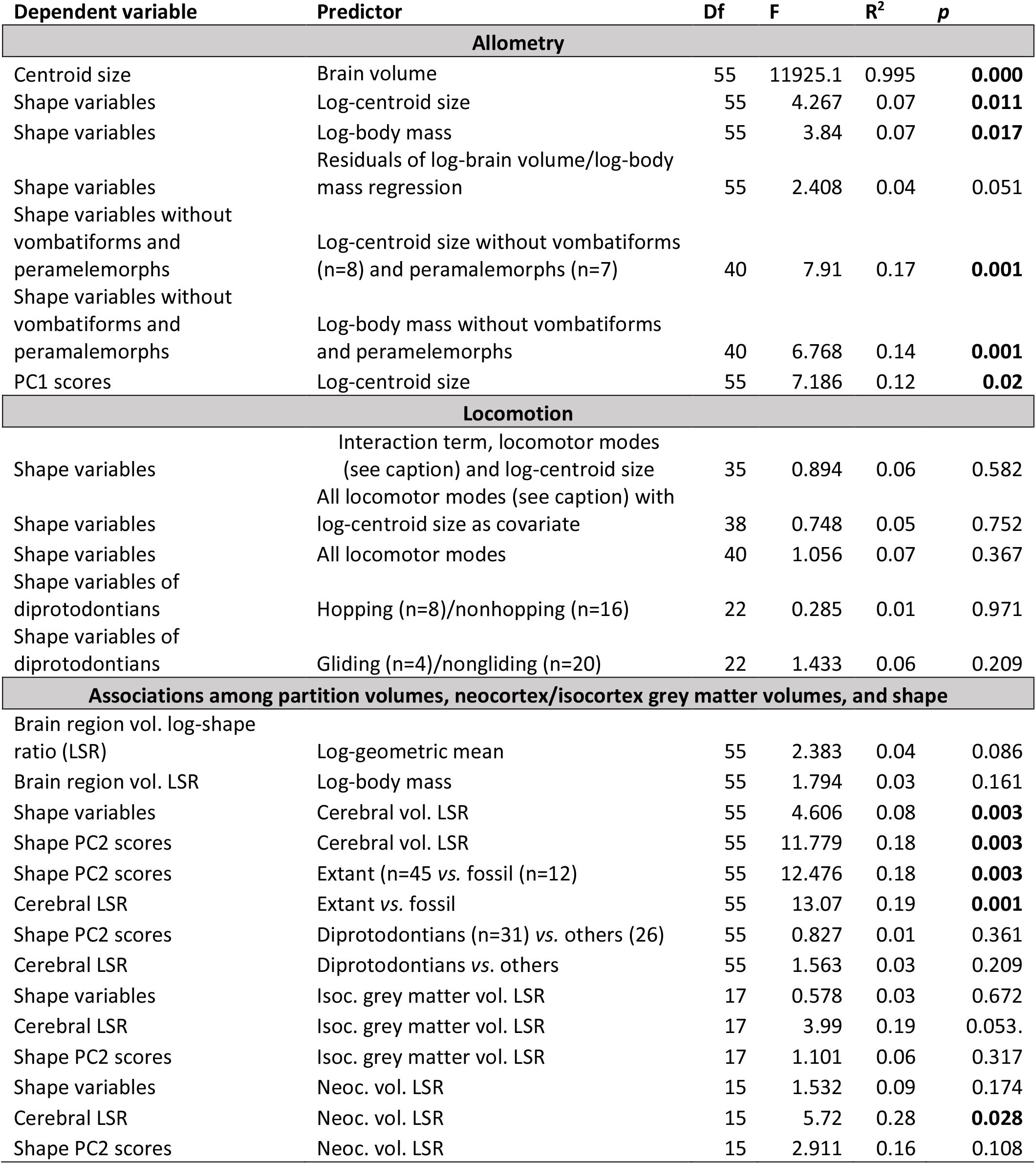
Summary table of all PGLS analyses conducted in this study. Df, degrees of freedom; F, F-value of effect size; R^2^, R-squared value; *p*, probability of no effect. Sample sizes for factorial categories are noted in the predictor column, except for arboreal (n=11+2 scansorial species) and terrestrial (n=19); locomotion. Bold *p* – values denote values below our significance threshold of 0.05.

| Dependent variable | Predictor | Df | F | $R^2$ | $p$ |
| --- | --- | --- | --- | --- | --- |
| **Allometry** |  |  |  |  |  |
| Centroid size | Brain volume | 55 | 11925.1 | 0.995 | **0.000** |
| Shape variables | Log-centroid size | 55 | 4.267 | 0.07 | **0.011** |
| Shape variables | Log-body mass | 55 | 3.84 | 0.07 | **0.017** |
| Shape variables | Residuals of log-brain volume/log-body mass regression | 55 | 2.408 | 0.04 | 0.051 |
| Shape variables without vombatiforms and peramelemorphs | Log-centroid size without vombatiforms (n=8) and peramelemorphs (n=7) | 40 | 7.91 | 0.17 | **0.001** |
| Shape variables without vombatiforms and peramelemorphs | Log-body mass without vombatiforms and peramelemorphs | 40 | 6.768 | 0.14 | **0.001** |
| PC1 scores | Log-centroid size | 55 | 7.186 | 0.12 | **0.02** |
| **Locomotion** |  |  |  |  |  |
| Shape variables | Interaction term, locomotor modes (see caption) and log-centroid size | 35 | 0.894 | 0.06 | 0.582 |
| Shape variables | All locomotor modes (see caption) with log-centroid size as covariate | 38 | 0.748 | 0.05 | 0.752 |
| Shape variables | All locomotor modes | 40 | 1.056 | 0.07 | 0.367 |
| Shape variables of diprotodontians | Hopping (n=8)/nonhopping (n=16) | 22 | 0.285 | 0.01 | 0.971 |
| Shape variables of diprotodontians | Gliding (n=4)/nongliding (n=20) | 22 | 1.433 | 0.06 | 0.209 |
| **Associations among partition volumes, neocortex/isocortex grey matter volumes, and shape** |  |  |  |  |  |
| Brain region vol. log-shape ratio (LSR) | Log-geometric mean | 55 | 2.383 | 0.04 | 0.086 |
| Brain region vol. LSR | Log-body mass | 55 | 1.794 | 0.03 | 0.161 |
| Shape variables | Cerebral vol. LSR | 55 | 4.606 | 0.08 | **0.003** |
| Shape PC2 scores | Cerebral vol. LSR | 55 | 11.779 | 0.18 | **0.003** |
| Shape PC2 scores | Extant (n=45 vs. fossil (n=12) | 55 | 12.476 | 0.18 | **0.003** |
| Cerebral LSR | Extant vs. fossil | 55 | 13.07 | 0.19 | **0.001** |
| Shape PC2 scores | Diprotodontians (n=31) vs. others (26) | 55 | 0.827 | 0.01 | 0.361 |
| Cerebral LSR | Diprotodontians vs. others | 55 | 1.563 | 0.03 | 0.209 |
| Shape variables | Isoc. grey matter vol. LSR | 17 | 0.578 | 0.03 | 0.672 |
| Cerebral LSR | Isoc. grey matter vol. LSR | 17 | 3.99 | 0.19 | 0.053. |
| Shape PC2 scores | Isoc. grey matter vol. LSR | 17 | 1.101 | 0.06 | 0.317 |
| Shape variables | Neoc. vol. LSR | 15 | 1.532 | 0.09 | 0.174 |
| Cerebral LSR | Neoc. vol. LSR | 15 | 5.72 | 0.28 | **0.028** |
| Shape PC2 scores | Neoc. vol. LSR | 15 | 2.911 | 0.16 | 0.108 |

We found no statistical association between any locomotor mode and brain shape (Table 1). However, locomotor mode is phylogenetically confounded among marsupials, particularly in relation to the single evolutionary transformation towards an upright posture related to hopping. This is relevant because the hopping kangaroos visually appear to score higher on PC2 than most other marsupials, except the possibly non-upright fossils *Ekaltadeta ima* and *Balbaroo nalima* (Den Boer et al. 2019), which are among the lowest-scoring diprotodontians (Fig. 3). However, we cannot distinguish whether this relates to locomotor mode or represents a random effect of Brownian motion. Note that differences between Macropodiformes and the remaining marsupials is concentrated in a dorsal and anterior expansion of the cerebral hemispheres, rather than the region of potential postural differences around the back of the brain (Russo and Kirk 2013), as shown in a follow-up visualisation comparing the mean landmark configurations of kangaroos *versus* other Diprodotontia (Supplementary Fig. 6). The arboreal dasyurid *Phascogale tapoatafa* and arboreal diprotodontians are widely separated in PC morphospace; similarly, the possum genera *Petaurus, Petauroides*, and *Acrobates*, which have independently evolved gliding adaptations, have widely diverging brain shapes in PC1/PC2 space. It is also noteworthy that the ecologically very similar wombat genera *Vombatus* and *Lasiorhinus* are widely separated on PC1.

### Intraspecific variation of endocast shape

A PCA of all specimens revealed substantial intraspecific variation in species represented by more than one specimen, with several instances where species overlap in PC morphospace but individuals of the same species are well separated (Fig. 5). This is consistent with clear visual differences of brain shapes within some species (see Supplementary Fig. 7, which compares strikingly different endocasts of two red kangaroo [*Macropus rufus*] individuals). However, comparison between intraspecific and interspecific Euclidean distances in Procrustes space (Supplementary Fig. 8) shows that, in terms of overall shape, intraspecific shape variation is much lower than interspecific variation.

**Figure 5:**
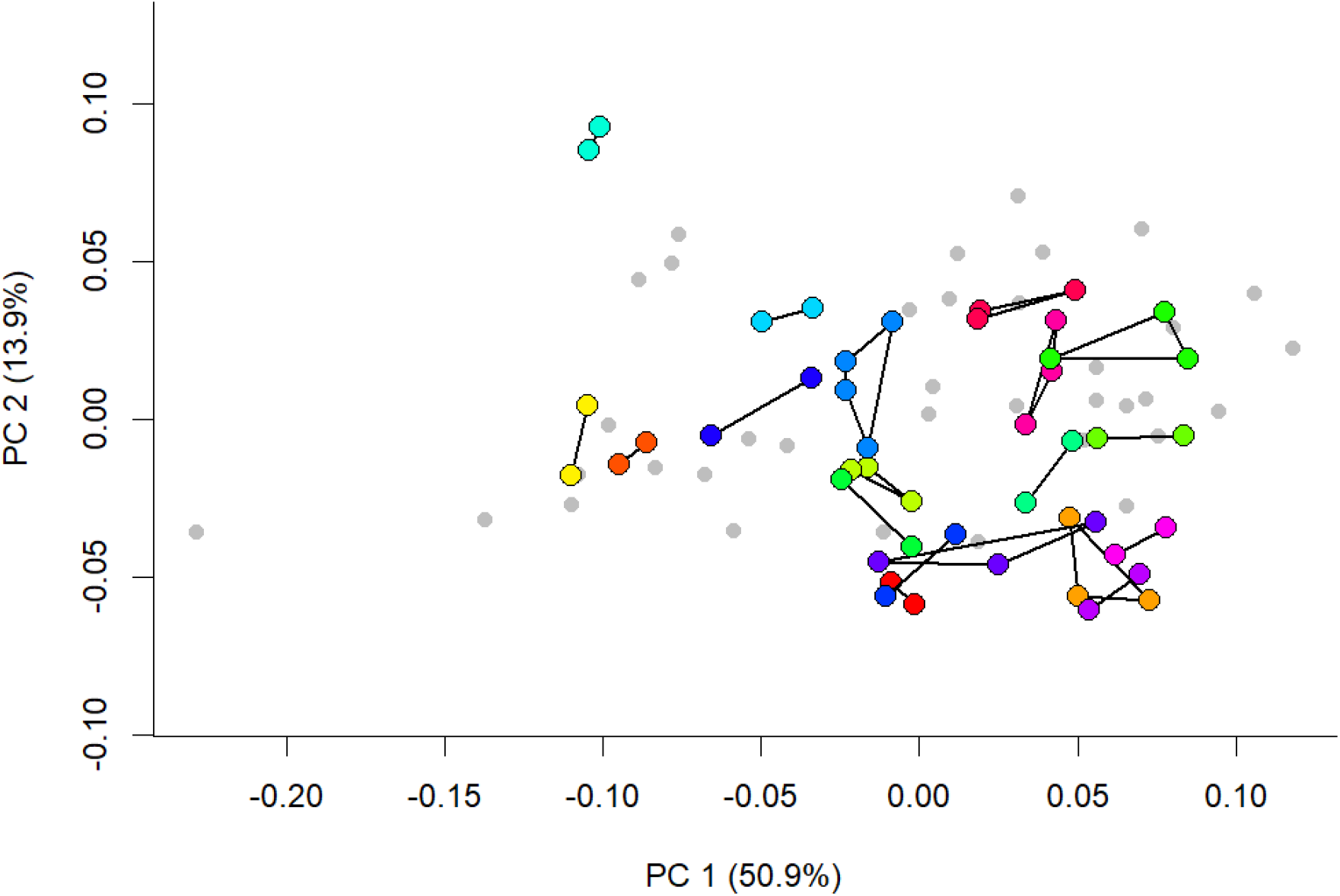
Plot of principal component 1 vs PC2 for all specimens according to their brain shape. Species with a single representative are grey points. Members of one species are coloured (arbitrarily) and connected by lines, demonstrating intraspecific variation in brain shapes along the two main axes of shape variation. Within-species distances compared to among species distances in Procrustes space are shown in Supplementary Fig. 8.

### Comparison of endocast shape with partition volume evolution, and relationship with histological neo/isocortical volumes

By comparison with endocast shape, the relative volumes of the brain partitions have slightly higher phylogenetic signal (K_mult_ = 0.66, p=0.001), but have no signal of evolutionary allometry (Table 1). A Mantel test of pairwise distances between species in the shape and volume PCA spaces is significant (*p*=0.000, but with low Pearson correlation values of 0.27), confirming our expectation that the two describe different patterns of evolutionary diversification (Fig. 6 shows the distribution of species on a volume-based PCA); for example, the PC1 of volumes shows high loading of olfactory bulb volume that is not apparent along the main variation of shapes (Fig. 6; supplementary movie 1). However, because PC2 of endocast shape appeared to reflect volumetric dominance of the cerebrum, and could possibly be associated with “neocorticalisation”, we ascertained that fossil species (n=12) indeed have significantly lower PC2 scores than extant species; fossil status explained nearly a fifth of PC2 (R^2^ =0.18; (Table 1)), but note that PC2 itself only explains 12.5% of endocast shape variation. We then assessed if relative cerebral size (as identified by the log shape ratio value of the cerebrum) was associated with endocast shape or PC2 of the shape PCA. This revealed a significant association of cerebral partition volume with shape and PC2 (Table 1), but neither association explained much variation. In keeping with their significantly smaller PC2 scores, fossil taxa also had significantly smaller cerebra relative to the remainder of brain partition volumes, and fossil status explained nearly 20% of variation in cerebral volume (Table 1). However, despite appearing to also have relatively larger PC2 scores and cerebral partition LSR, diprotodontians did not have significantly higher PC2 scores or Cerebral LSR in a phylogenetic context (Table 1).

**Figure 6:**
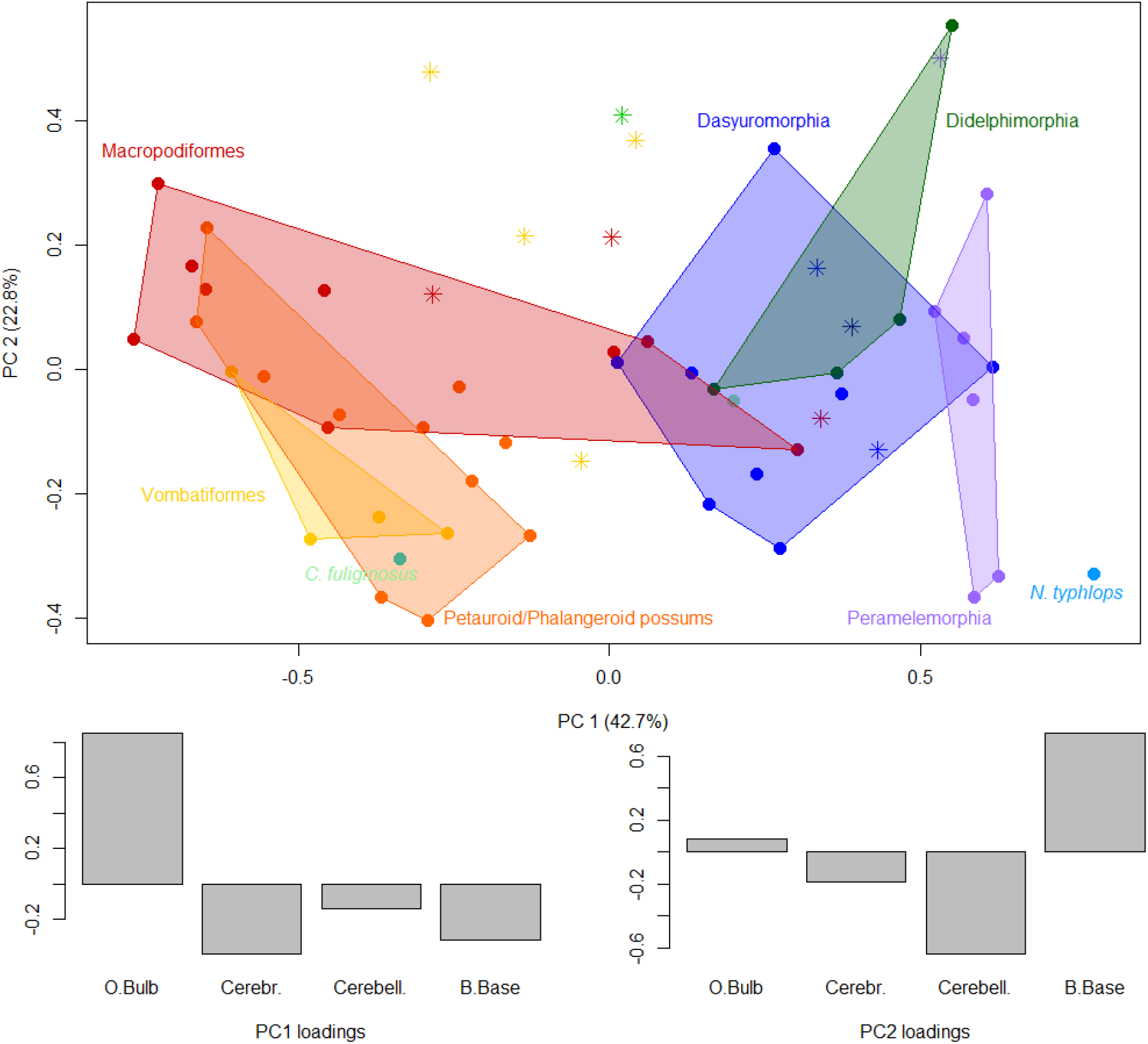
Plot of principal component 1 vs PC2 of species according to their brain region volume (top) and PC1 and PC2 variable loadings (bottom) based on log-shape ratios of four endocast partitions: olfactory bulb [O.bulb], cerebrum [Cerebr.], cerebellum [Cerebell.] and brain base [B.Base]. Note that the main differentiation between species here is due to volumes of the olfactory bulb. Colouration as in Fig. 1 and 2.

We found no dependable association between shape variation, relative volume variation, and neocortical or isocortical volume. The only significant association we found was a small association between the log-shape ratios of cerebral hemispheres and neocortex volumes (across 17 species) (Table 1). Therefore, although cerebral hemisphere partition volume may be indicative of neocortex volume, this association is not strong and cannot be translated to shape variation.

## Discussion

Our results reveal that over half of marsupial endocast shape variation lies along a spectrum from elongate/straight to globular/inclined endocast shapes, which strongly resemble the “spatial packing” effects proposed for primates and experiments on mice (Ross and Henneberg 1995; Lieberman et al. 2008; Bastir et al. 2011; Marcucio et al. 2011; Zollikofer et al. 2017). However, “spatial packing” was suggested to accommodate relatively larger brains into skulls with short cranial bases, while the marsupial pattern is not associated with relative brain size. The spectrum of shapes is also very broad, ranging from nearly spherical in the marsupial mole *Notoryctes typhlops* to nearly tubular in species like the fossil vombatiform *Silvabestius johnnilandi* (compare on Fig. 1). This resembles other instances where an axis of global change related to elongation determines morphological diversification, for example in the whole body of fishes, lizards, and mustelids (Bergmann and Irschick 2010; Ward and Mehta 2010; Law et al. 2019). Shape diversification partially associated with elongation has also been postulated for the therian cranium (e.g. Cardini et al. 2015), although this so-called “Cranial Rule of Evolutionary Allometry” is also associated with size variation, which has no strong influence in our dataset.

The high variability of endocast shapes across short evolutionary time scales (or even within species) suggests that the diverse brain tissues within the endocast are integrated to produce a limited set of shapes reflected in PC1 (Felice et al. 2018). But what determines this strong main axis of variation? The pattern resembles an evolutionary “line of least resistance” (Schluter 1996), a genetically favoured direction of morphological evolution. In mammals, strong allometric variation is often portrayed as a hallmark of such evolutionary lines of least resistance (e.g. Marroig and Cheverud 2005; Marcy et al. 2020). However, the pattern we observe is not necessarily genetic and shows no strong allometry. In addition, endocasts differ from whole-body or overall cranial shape through the juncture between the soft tissue of the brain with its neuronal functionality on one hand, and the osseous skull with its functional diversity on the other.

Focusing on the juncture between brain and skull tissue, brain shape is known to be mechanically malleable. For example, it can be determined by intrinsic mechanical properties (such as tissue stiffness or internal tension of neurons; Atkinson et al. 2015; Koser et al. 2016; Heuer and Toro 2019), as well as external impacts (e.g. contact with cranial bones during brain growth; Macrini et al. 2007c; Budday et al. 2015). Such impacts are known to change brain shape over short evolutionary time scales and also individual development (Budday et al. 2015; Gómez-Robles et al. 2015), just as observed here. Similarly, brain sizes and regional volumes (and therefore presumably their shape) can vary substantially during individual lifetimes (Burger et al. 2013; Dechmann et al. 2017). Thus, mechanistic processes may mould brains into default shapes which we see on PC1, but without representing a “true” constraint because the shape of the brain is not intrinsically fixed. This is also consistent with our observations of highly unique endocranial shapes. Particularly striking examples of this are the deep “waist” between the cerebrum and the cerebellum, reflecting a deep imprint of the highly pneumatised mastoid part of the petrosal of the pygmy glider *Acrobates pygmaeus*, or the flat brain with imprint of the middle ear cavity in *Planigale ingrami* (compare in Fig. 1)

With respect to possible associations between brain shape and skull functionality, neither allometry or locomotor mode were strongly associated with endocast shape. There are at least two functional reasons to explain this: First, smaller mammals have relatively larger brains for their body mass, and would therefore be expected to share common constraints of distributing a heavy brain in a small skull. In addition, as postulated under the “spatial packing” hypothesis, spatial constraints might require brains to be packed in a more globular way when their mass increases relative to body mass. However, allometry is not an important part of brain shape variation (mirroring results from a study on squirrels and their fossil relatives; Bertrand et al. 2019b) and our “spatial packing” - like pattern exists without allometry; the much higher phylogenetic signal of centroid size compared to endocast shape also demonstrates the relative evolutionary independence of the two. We also found no association of locomotor categories with endocast shape, including a lack of association between gliding and endocast shape, which contrasts with findings in gliding squirrels (Bertrand et al. 2019b). There is also no obvious connection between facial length and brain globularity (Evans et al. 2017; Zollikofer et al. 2017): for example, the two most globular brains in the sample belong to the relatively short-snouted marsupial mole and the relatively long-snouted shrew opossum (see Fig. 2).

The lack of functionally explainable pattern of endocast shape can have several methodological causes related to a relatively broad landmarking protocol, broad locomotor categories, and the phylogenetically confounded distribution of locomotor mode (for successful, phylogenetically more restricted analyses of ecology and endocast shape, see Ahrens 2014; Bertrand et al. 2019b). However, it is also possible that the evolutionary flexibility of brain shape suggested above, and its apparent propensity to adapt to the braincase, might result in noisy and divergent adaptations that are statistically not tractable. This also emphasises that interpreting associations between locomotor mode and endocast shape, as well as possibly many other adaptations such as feeding biology, require consideration of whether variation is caused by cranial, rather than somatosensory, adaptation (Jeffery and Spoor 2006). However, it is also possible that the skull is not always a strong predictor of locomotor mode; for example, the skull shape of kangaroos is not strongly associated with hopping or different modes of hopping (Butler et al. 2020), possibly explaining why we also found no significant association between endocasts of hopping *vs*. non-hopping kangaroos.

We find that different descriptions of endocast or brain macromorphology (shape coordinates, volumetric dissection, and functional region volume such as neocortex) are not interchangeable in terms of the information they contain. What little significant correspondence we find (e.g. that overall shape, PC2, and neocortex volume are significantly associated with relative cerebral hemisphere size) is not sufficient to allow biological conclusions from one measure to the other. This also appears to be the case with specific somatosensory regions such as the visual cortex, which occupies the region that expands most along PC2 (Karlen and Krubitzer 2007). An association between occipital expansion of the neocortex and a potential improvement of vision capabilities has been suggested in rodents and their relatives (e.g. Bertrand et al. 2019a). In marsupials, however, a previous study (Karlen and Krubitzer 2007) showed no differences in visual cortex area between two species with high PC2 scores (*Dactylopsila trivirgata* and *Trichosurus vulpecula*) *versus* species with low PC2 scores (*Dasyurus hallucatus, Monodelphis domestica*, and *Didelphis virginiana*). In addition, the marsupial mole *Notoryctes typhlops* with its dorsally rounded cerebral hemispheres (Fig. 1) has no functional eyes (Van Dyck and Strahan 2008).

Fossil species have significantly smaller relative cerebral size and PC2 scores than their living relatives, which superficially supports the notion of increasing neocortex dominance (“neocorticalisation”) in mammals (Jerison 2012), and specifically marsupial (Haight and Murray 1981), evolution. However, very weak associations between shape, cerebral, and histological cortical volumes suggest that surface area measurements (Jerison 1973; Bertrand et al. 2016) on a sample where these measurements can be made with confidence will be required to clarify this effect.

Phylogenetic divisions in brain shape have been successfully established at finer phylogenetic scales (e.g. Silcox et al. 2009; Thiery and Ducrocq 2015; Bertrand et al. 2016; Bertrand et al. 2019b). We also find moderate phylogenetic signal in brain shape. However, this differentiation is concentrated on the second principal component and explains little shape variation (compare the mean shapes with the shapes of individual species in Fig. 1 or supplementary Figure 5). Brain shape is, therefore, likely too ambiguous to be useful in marsupial phylogenetics, with anatomical scores that reflect the internal bony surface likely more successful (Haight and Murray 1981; Macrini et al. 2007b).

## Conclusions

High evolutionary shape plasticity along a global axis of elongation emerges as a powerful mechanism of balancing the evolution of marsupial, and possibly mammalian, cranial function against the need to accommodate the brain. However, the precise mechanisms for this flexibility remain to be understood and are likely quite diverse. For example, on one hand, brain shape and volume proportions can undergo drastic seasonal change in some small mammals (Dechmann et al. 2017). On the other hand, dogs bred for specific, small cranial vault shapes can display pathological cerebellar compression (Hechler and Moore 2018). Lastly, over the large time scales of mammalian brain evolution, increases in brain size appear to have triggered cranial vault expansion through a heterochronic delay in ossification of the cranial roof (Koyabu et al. 2014). The main challenge for understanding how the mammalian brain co-evolves with the skull will therefore be in separating the effects of individual developmental flexibility of brain shape on one hand, and deep time co-evolution between the cranial vault and brain size on the other, in a broader sample of non-primate mammals.

## Author contributions

VW conceived of the study, performed most of the endocasting, co-analysed the data, and wrote the manuscript. TR, SW, TEM, KJT, KB, MA, SH, JB, RMDB, SL, AS, provided specimen scans, co-wrote the manuscript, and co-developed the palaeontological aspects of the study. RMDB produced the phylogenies. KG participated in endocasting and most of the endocast dissections, and co-developed the dissection protocol. KM scanned a proportion of specimens, and co-wrote the manuscript. ES developed most R code, co-analysed the data, and co-wrote the manuscript.

## Acknowledgements

This research was funded by an Australian Research Council Discovery Early Career Award (DE120102034), Discovery Project DP170103227, and Future Fellowship FT180100634 to V.W; ARC DP170101420 to MA and SJH; ATM MNHN ‘Biodiversité actuelle et fossile’ to S.L.; and a University of Adelaide Fellowship to E.S. Scans provided by the Digimorph platform were funded by National Science Foundation (NSF) IIS 9874781 and IIS 0208675 (TR), and several scans were done by T.M. under NSF DEB 9873663 (Nancy Simmons, AMNH) and NSF DEB 0309369 (Rowe and Macrini). For involvement in the collection of the fossils, we thank P. Creaser and the CREATE Fund, K. and M. Pettit, the University of New South Wales, Environment Australia, Queensland Parks and Wildlife Service, Outback at Isa, Mount Isa City Council, The Riversleigh Society and the Waanyi people. We thank the late Simon Collins (School Veterinary Science, UWQ) and Stephen Johnston (School of Agricultural and food Sciences, UQ) for making available CT scans of two *Phascolarctos cinereus*. We thank Candi Wong and Narelle Hill for support with endocast segmentation and dissection. The authors acknowledge the facilities and scientific and technical assistance of the National Imaging Facility, a National Collaborative Research Infrastructure Strategy (NCRIS) capability, at the Centre for Advanced Imaging, University of Queensland. We thank Materialise Malaysia for assistance with endocast preparation in Mimics. This study was mostly conducted on the traditional lands of the Yuggera (Brisbane) and Kaurna (Adelaide) people.

## Data Availability

All endocasts are available on Figshare: For the original stl files and endocast dissections, the doi is http://10.0.23.196/m9.figshare.12284456). For ply files used for automatic landmark placement, the doi is http://10.0.23.196/m9.figshare.12284456. The full R code for data read-in, analysis, and Figures, including, auxiliary files, volume data, and landmark coordinate data are available in github repository https://github.com/VWeisbecker/Brain_shape_study (see README instructions there).

---

**Supplementary Figure 1:**
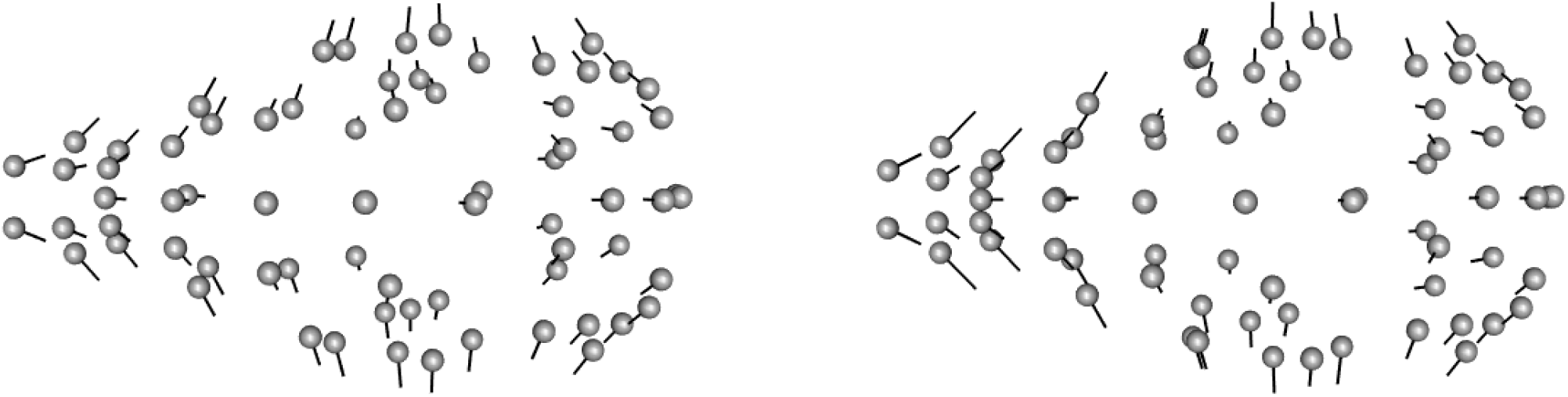
Landmark displacement graphs of ordinated endocast shape for landmark coordinates with surface semilandmarks automatically placed (left) and manually placed (right) along the first Principal Component (PC1), showing the displacement of landmarks from the extreme shape associated with PC1 minimum scores (grey spheres) to the mean shape (end of hairlines). This comparison is on the full dataset without symmetry removal or averaging of specimens within a species.

**Supplementary Figure 2:**
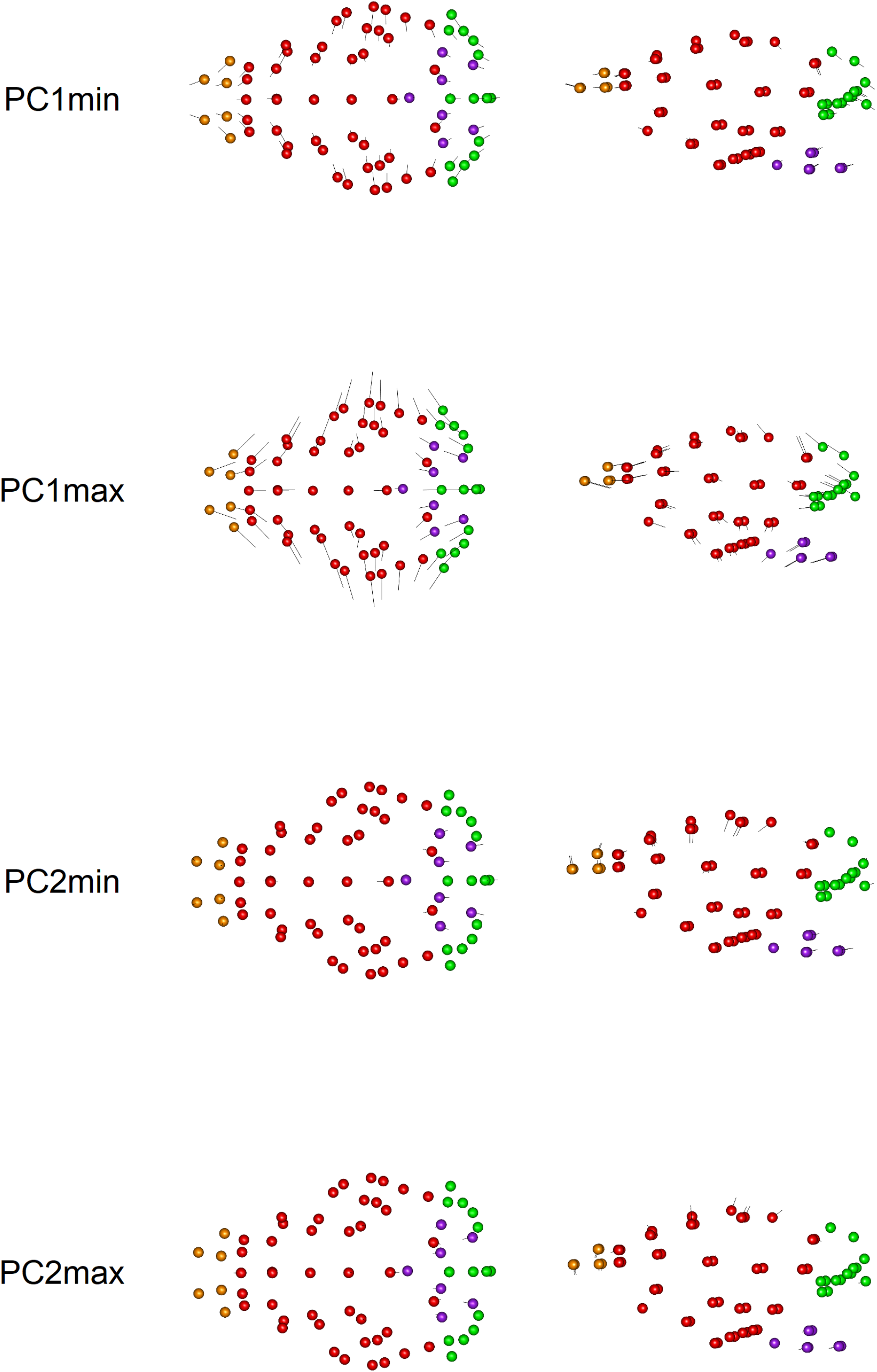
Landmark displacement graphs of shape variation along PC1 and PC2, with the mean shape as reference (spheres) and the “hairs” of the vectors pointing to Principal Components score extremes for PC1 and PC2.

**Supplementary Figure 3.**
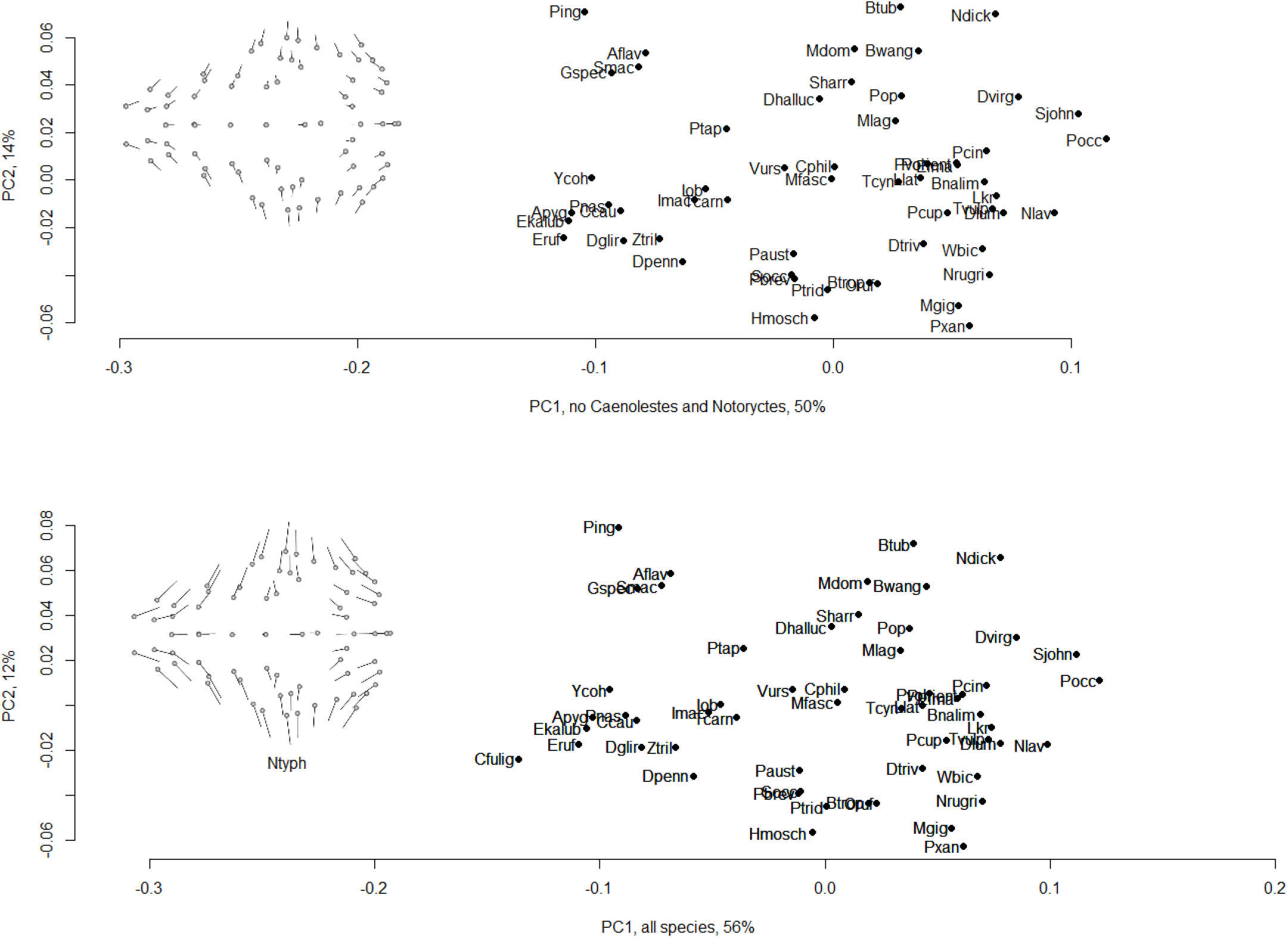
Principal Components (PC) plots of species with (bottom) and without (top) the two highly spherical endocasts of the marsupial mole, *Notoryctes typhlops*, and the shrew-opossum, *Caenolestes fuliginosus*. Note that the removal of these two species caused species to have very similar relative positions, but switched PC1 score signs; the signs of the PC1 scores in the plot including all species (bottom) were therefore also switched to facilitate comparison.

**Supplementary Figure 4:**
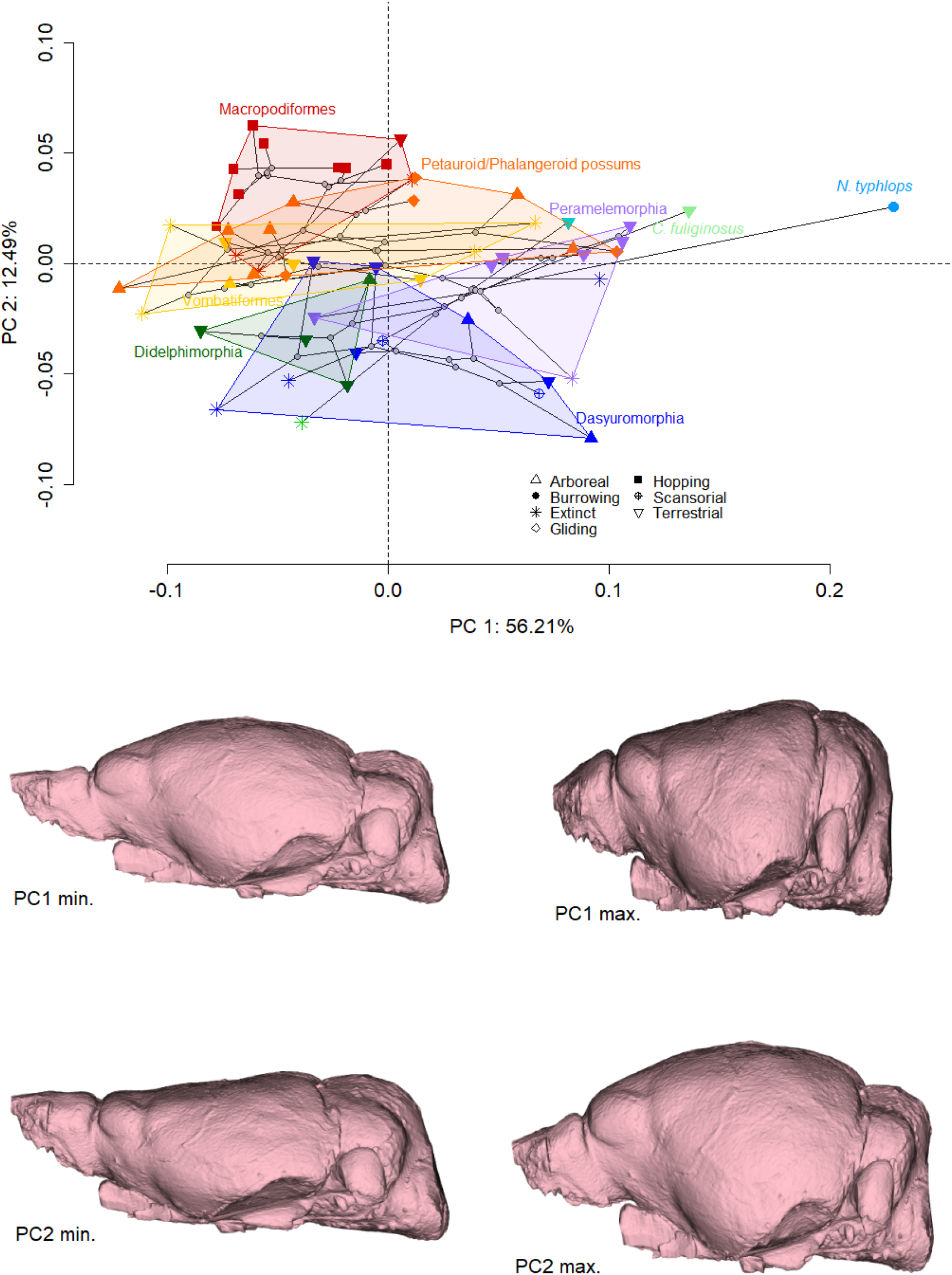
Plot of principal component 1 vs PC2 for 57 species of marsupials according to their brain shape (top) and corresponding shape variation illustrated as warped endocasts representing the highest and lowest PC1 and 2 scores (bottom). Polygons are drawn around living members of all major clades and their extinct relatives. Stars represent extinct species.

**Supplementary Figure 5:**
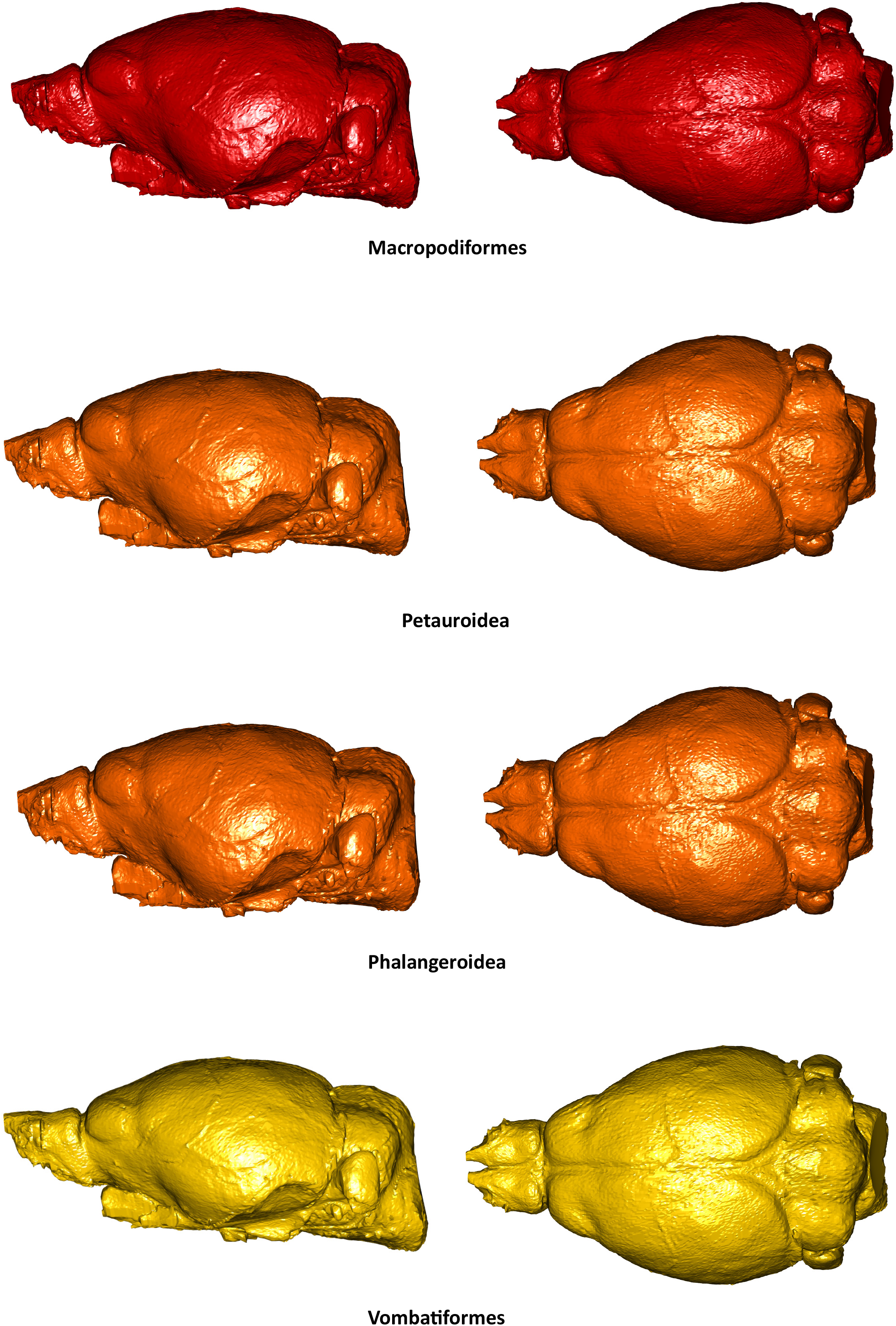
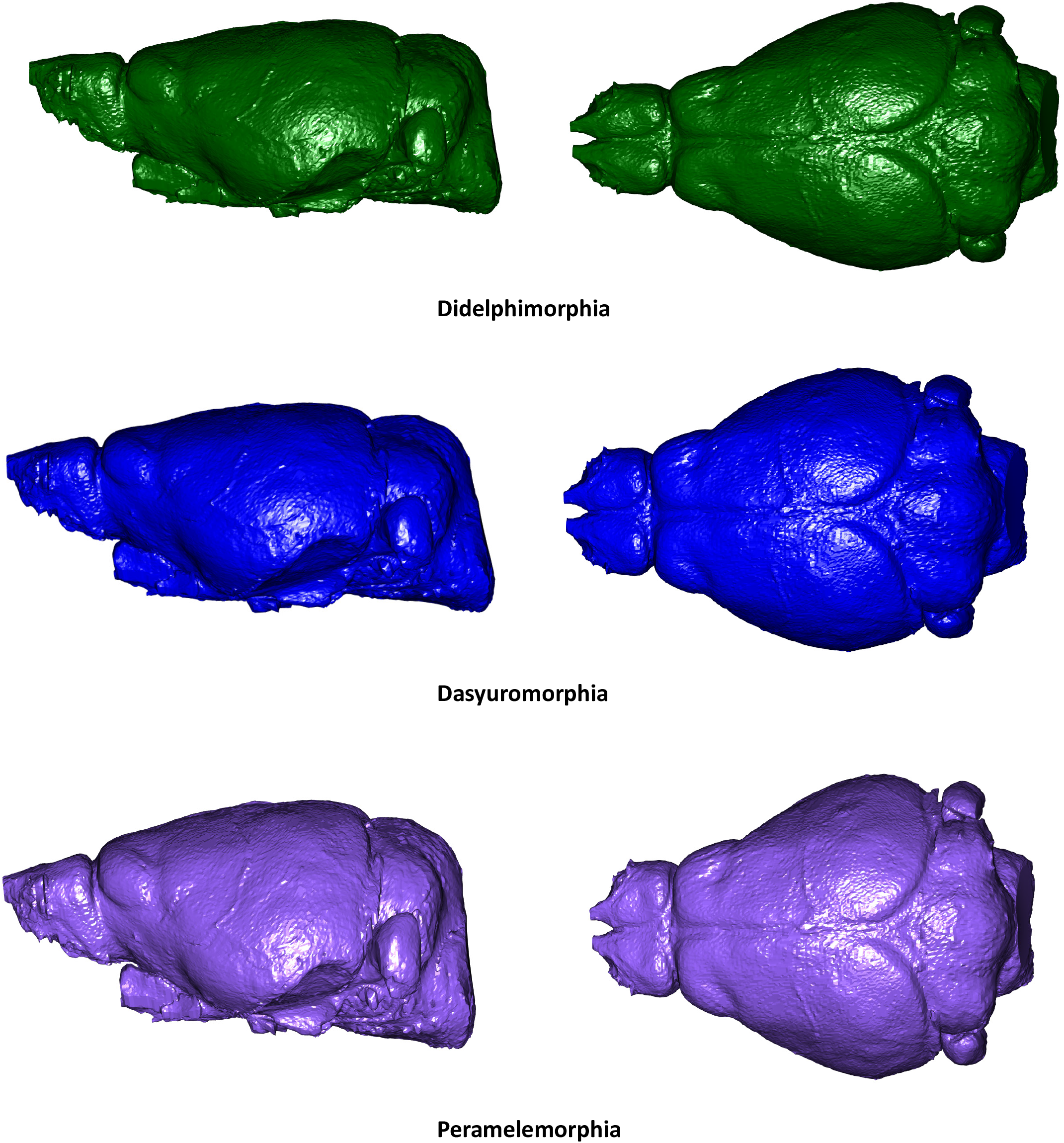
Mean shapes for the Diprotodontian major clades and the three remaining orders of marsupials for which multiple speciems were available.

**Supplementary Figure 6:**
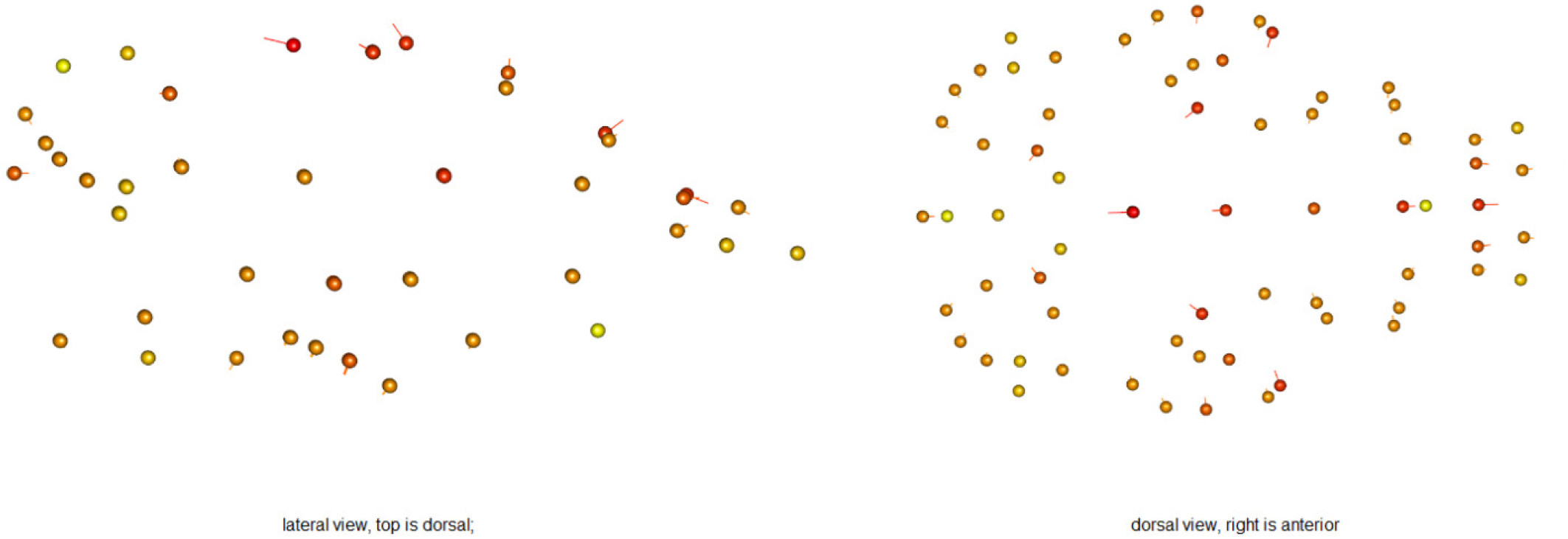
Lateral (left) and dorsal (right) heat plots of the distribution of shape variation magnitudes in the mean configuration of diprotodontial marsupials (balls) compared to the mean landmark configuration of macropodiforms (kangaroos; ends of hairline vectors). The colouration is calibrated for the greatest magnitude in landmark displacement to be darkest red, and the least magnitude to be the lightest yellow. Note the lack of substantial movement around the brain base.

**Supplementary Figure 7:**
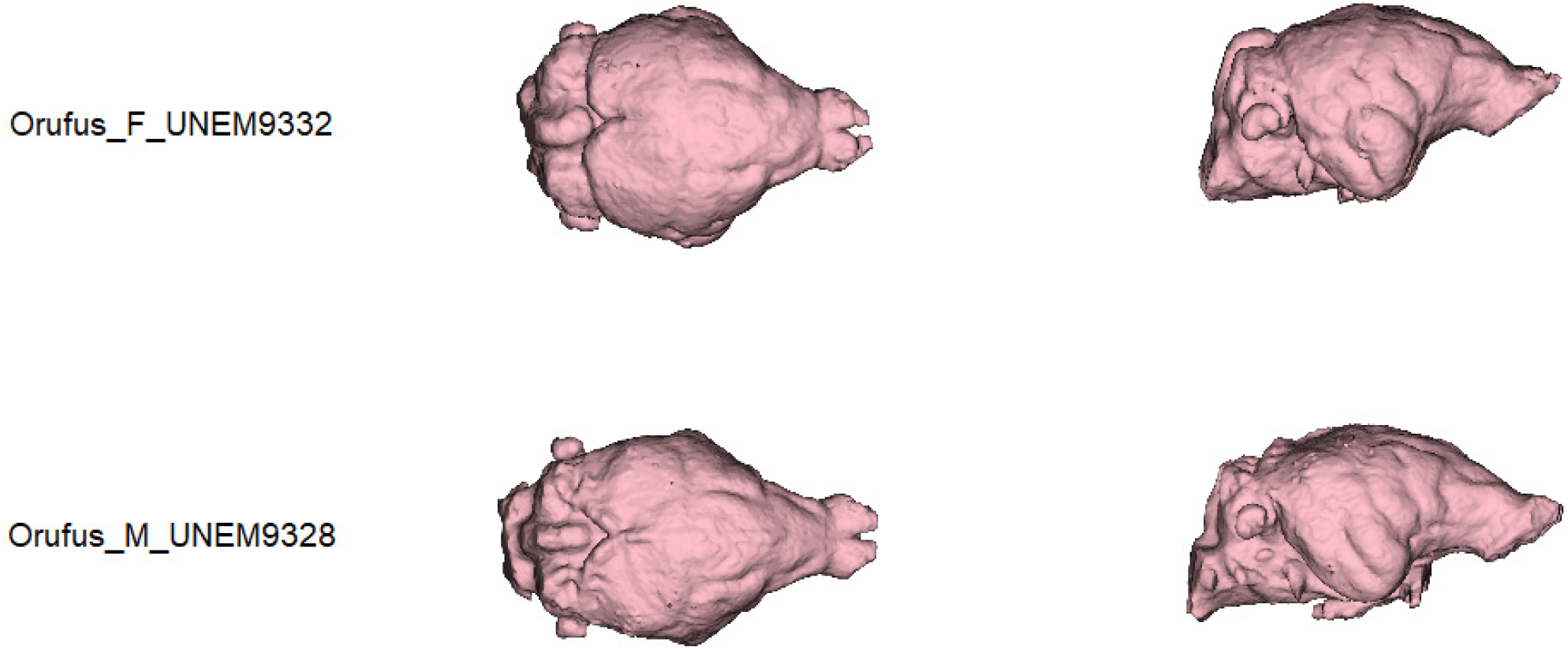
Dorsal (left) and lateral (right) images of brains of a female (top) and male (bottom) red kangaroo (*Osphranter rufus*). Note the substantial differences in shape, particularly in terms of width and inclination.

**Supplementary Figure 8:**
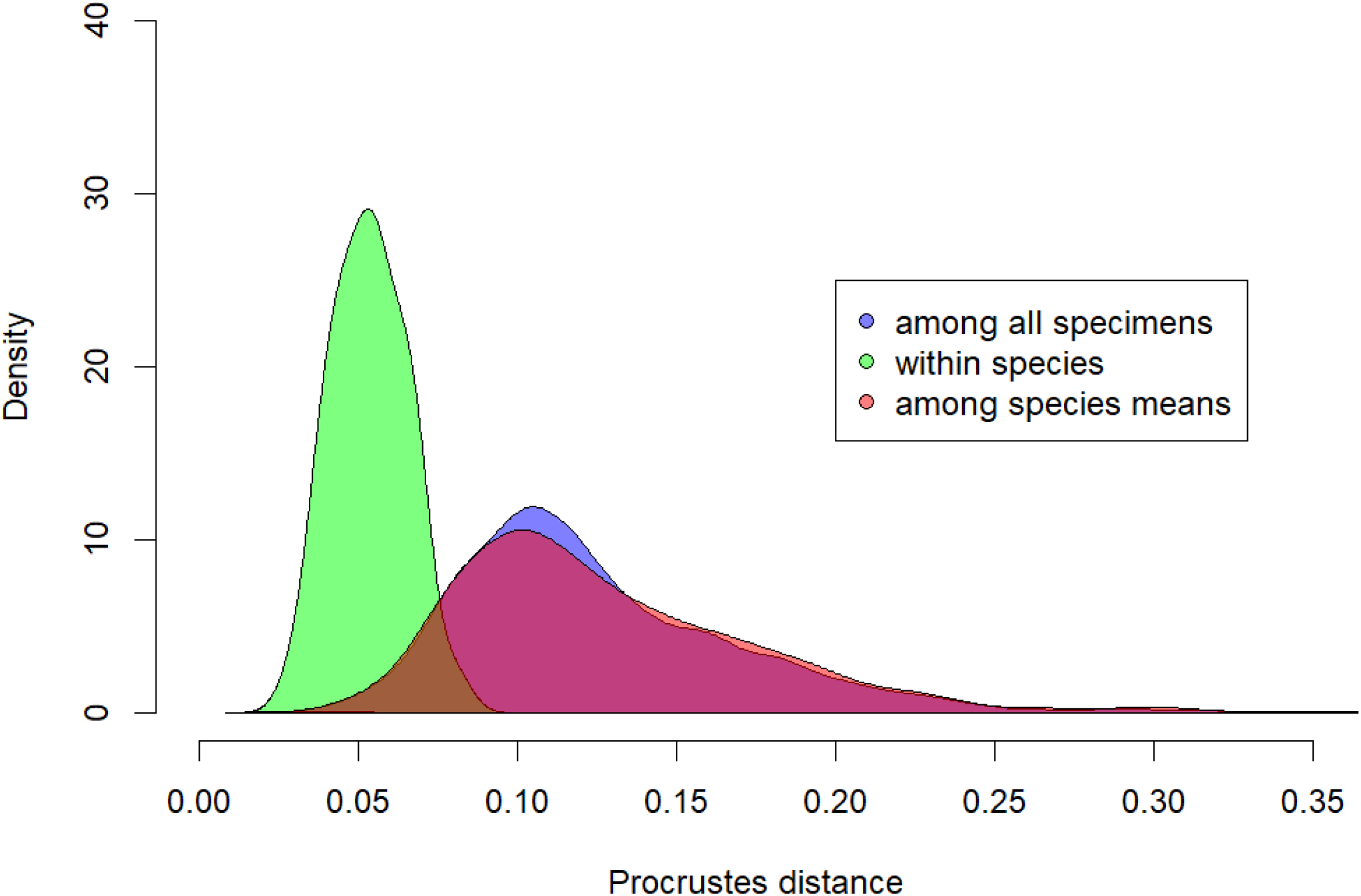
Density plots of the frequency of Procrustes distances (i.e. taking all of shape into consideration), showing that Procrustes distances within species are generally lower than among species means or among all specimens.

## Supplementary Information 1 Methodological details for landmark placement, endocast dissection, and the phylogenies used

### Landmarks

Refer to Fig. 1 below for illustration of landmarks. The numbers indicated below are the automatic patch placement; note that the procedure of automatic patch placement changes the landmark numbers from the numbers originally given in Checkpoint. The anatomical terminology was taken from Macrini *et al*. (2007).

#### Fixed landmarks

1. Left paraflocculus
2. Right paraflocculus
3. Anterior midpoint of hind brain
4. Base of olfactory bulb
5. Left Tip of olfactory bulb
6. Right Tip of olfactory bulb
7. Left hypoglossal foramen
8. Right hypoglossal foramen

#### Fixed landmarks of patch

The patch was constructed in checkpoint, which only produces square patches which are difficult to place on round objects like the brain. To enable the characterisation of the lateral sides of the brain, we therefore placed two-point curves in checkpoint between landmarks 25-22, 44-22, 47-24 (left) and 26-24, 50-22, 53-22 (right). However, landmarks shared with the patch and the curve landmarks thus obtained were slid as a patch, as seen in Fig. 2.

36: Point of divergence between the two cerebral hemispheres, just anterior to the transverse sinus
35,39: Center of indentation between cerebrum, lateral, and middle cerebellar lobe
21,23: Point on cerebrum anterior to prootic vein exit
37: Center of patch – on the cerebral midline and equidistant from points 38 and 36
25,26: lateral extremity of junction between olfactory bulb and cerebral hemispheres
38: midpoint of the cross described by the midline of the brain and the junction between olfactory bulb and cerebral hemispheres
21-22, 23-24: Point on cerebrum anterior to prootic vein exit (shared with patch) to point of divergence between cerebral hemisphere and emergence of V2 (maxillary branch of trigeminus)

#### Curves (all with two end points and two semilandmarks)

9-12: Base to top of middle cerebellar lobe
13-16, 17-20: posteriormost point of lateral cerebral lobe to highest point of parafloccular stalk
30-27: Connecting the lateral-most points of each olfactory bulb side.
35-39: connecting the ventral stalk nerves exiting the jugular foramen.

**Figure 1:**
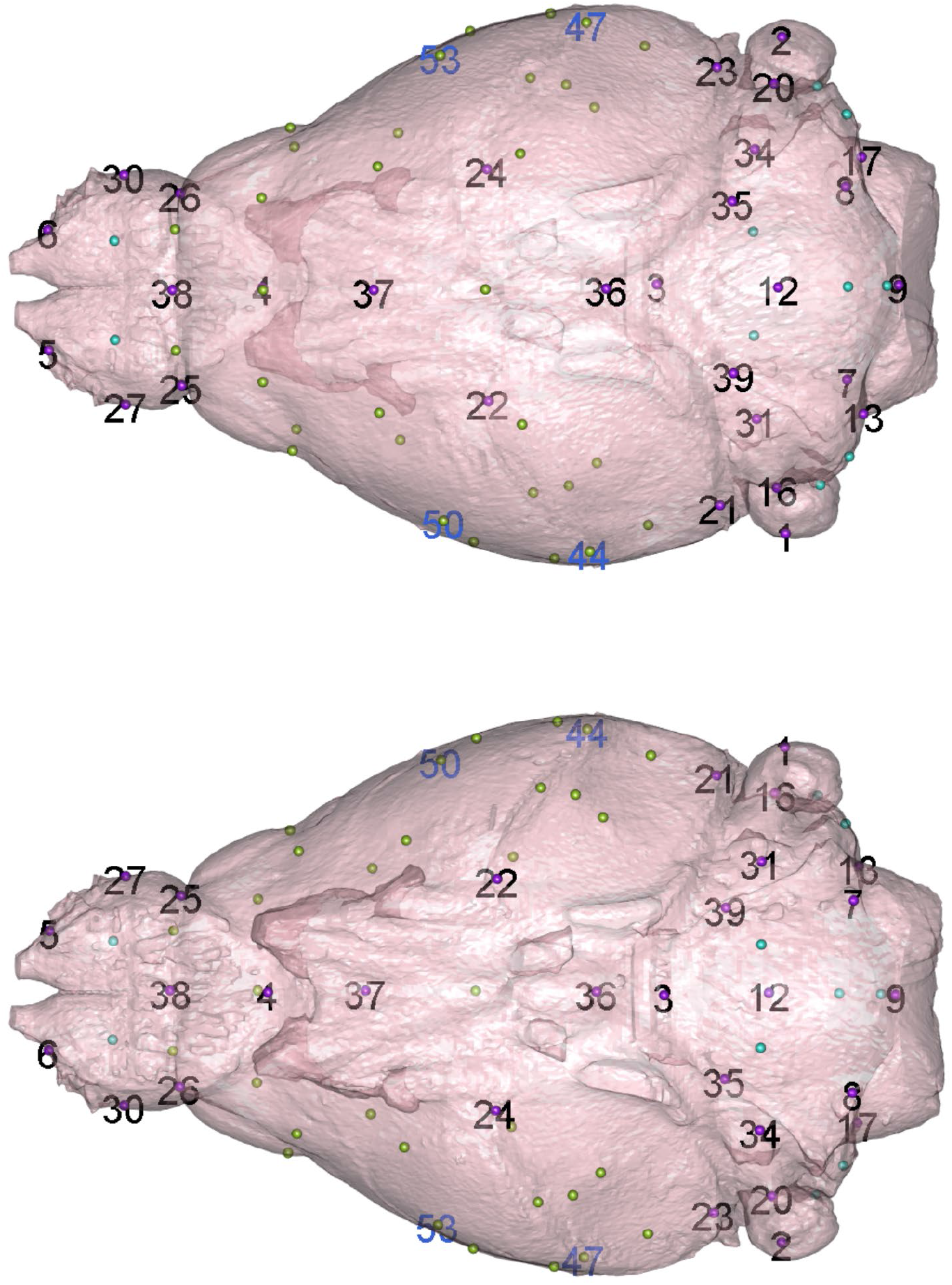
Numbers of fixed landmarks outlined above on the brain of *Phascogale tapoatafa* JM12395 (the specimen used to produce the atlas for automatic landmark placements), warped to the mean shape in geomorph. The blue numbers represent auxiliary curve origins placed in checkpoint to allow for the application of sliding surface semilandmarks on the lateral side of the brain. Numbers are given for all fixed landmarks, superimposed on the landmarks which are coloured as in Fig. 1.

### Endocast dissection

All sections were performed in Mimics V. 17-20, or for corrections of irregular cuts in 3Matics v. 9. See Fig. 2 for an illustration of the dissections. The dissected volumes are all available on Figshare repository http://10.0.23.196/m9.figshare.12284456

#### Olfactory bulb

Cut along the groove between the olfactory bulb and the cerebral cortex.

#### Cerebellum

Dorsal line along the junction between cerebrum and cerebellum; cut between vestibulo-cochlear nerve and paraflocculus; cut along the boundary of cerebellum and medulla.

#### Brain stem

Cut between vestibulo-cochlear nerve and paraflocculus; cut across the opening of the foramen magnum; cut across the medulla/pons boundary at the posterior end of the cerebral hemisphere behind the base of the foramen ovale of the trigeminal nerve (V5)

#### Cerebrum

Subtract the above three from the brain for cerebral volume

**Figure 2:**
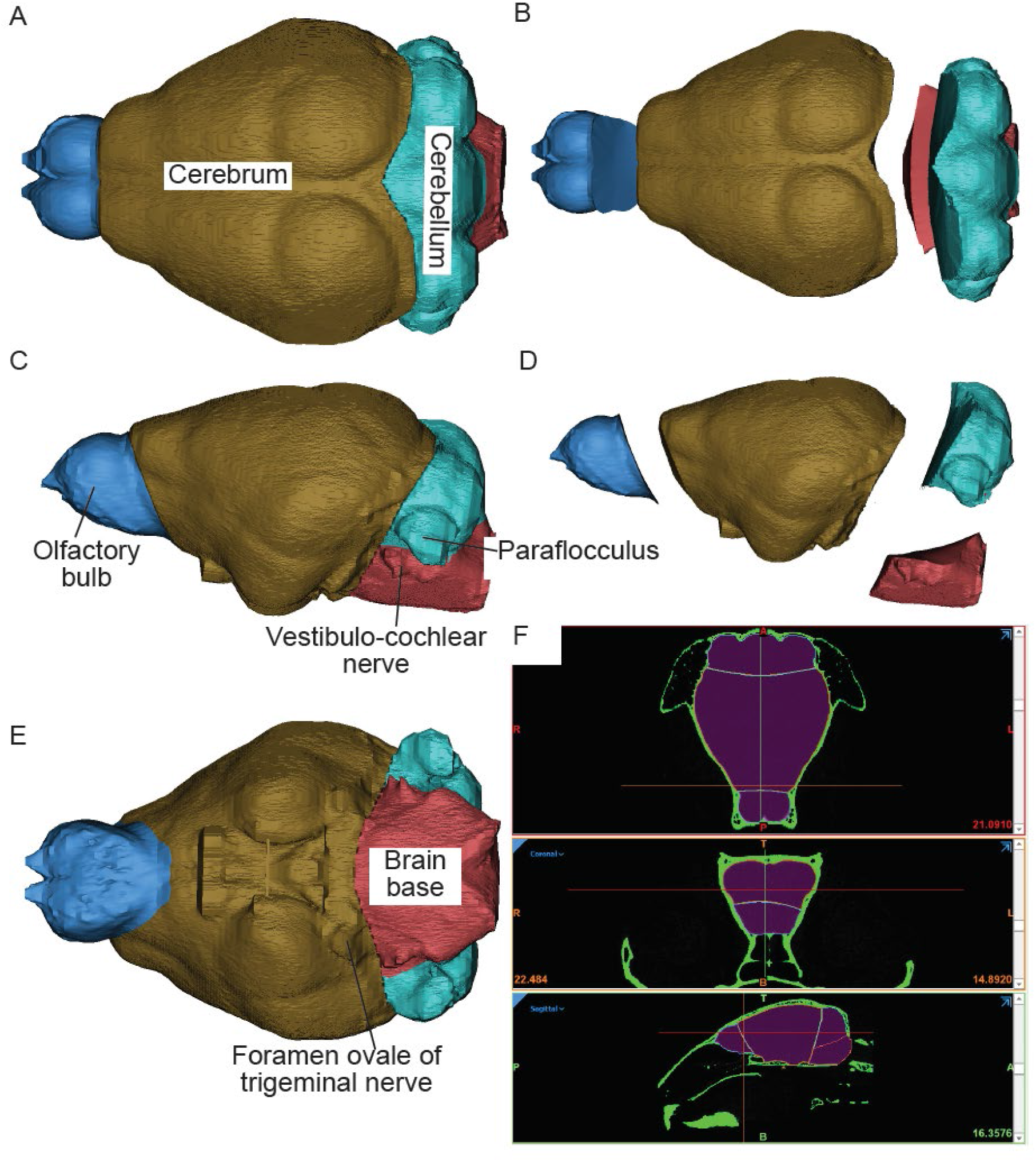
Details of endocast dissections in dorsal (A,B), lateral (C, D), and ventral (E) positions, illustrated on the brain of the sugar glider (Petaurus breviceps) TMMM82261. F) is the transverse, coronal, and sagittal view of the skull (in green) with the endocast in it (in pink). The thin lines reflect the lines along which the endocast was dissected.

### Phylogeny

Tree topology and divergence dates for living taxa follow Mitchell et al. (2014), except that the tree root was positioned between Didelphimorphia and the remaining extant orders following Gallus et al. (2015). *Borhyaena tuberata* is a member of Sparassodonta, which falls outside crown-clade Marsupialia(Forasiepi 2009); we have arbitrarily set the divergence of *Borhyaena* as 5 million years before the estimated time of origin of crown-clade Marsupialia. The macropodiforms *Balbaroo nalima* and *Ekaltadeta ima* are placed in a polytomy with Hypsiprymnodontidae, Potoroidae and Macropodidae (Black et al. 2014). The macropodid *Simosthenurus occidentalis* is assumed to have diverged at the same time as *Lagostrophus fasciatus* (Llamas et al. 2015). The fossil dasyuromorphian *Barinya wangala* is placed in a polytomy with Dasyuridae, Myrmecobiidae and Thylacinidae(Archer et al. Submitted), whereas the fossil thylacinid *Nimbacinus dicksoni* is assumed to have diverged from the recently extinct *Thylacinus* midway between the age of *Nimbacinus* (Woodhead et al. 2016) and the time of the Dasyuridae-Myrmecobiidae-Thylacinidae split. The extinct peramelemorphian *Galadi speciosus* is assumed to have diverged from the midpoint of the branch leading to crown-Peramelemorphia (Travouillon et al. 2015). Divergence dates for fossil vombatiforms are based on *Beck et al*. (Beck et al. 2014a). Three different positions for *Yalkaparidon coheni* were tested (following the phylogenetic analyses of Beck *et al*.(2014b)): stem-diprotodontian (diverging at the midpoint of the branch leading to crown-Diprotodontia), stem-australidelphian (diverging at the midpoint of the branch leading to crown-Australidelphia), and member of Agreodontia sensu Beck *et al*. (2014b) in a polytomy with Dasyuromorphia, Peramelemorphia and Notoryctemorphia. For all phylogenetic analyses, all three placements for *Yalkaparidon* were tested and results were averaged.

